# Unusual 1-3 peptidoglycan cross-links in *Acetobacteraceae* are made by L,D-transpeptidases with a catalytic domain distantly related to YkuD domains

**DOI:** 10.1101/2023.10.31.563487

**Authors:** Marcel G Alaman-Zarate, Brooks J Rady, Caroline A Evans, Brooke Pian, Darren Greetham, Sabrina Marecos-Ortiz, Mark J Dickman, Ian DEA Lidbury, Andrew L Lovering, Buz M Barstow, Stéphane Mesnage

**Author notes:** These authors contributed equally to this work (Marcel G Alaman-Zarate, Brooks J Rady).

## Abstract

Peptidoglycan is an essential component of the bacterial cell envelope that contains glycan chains substituted by short peptide stems. Peptide stems are polymerized by D,D-transpeptidases, which make bonds between the amino acid in position 4 of a donor stem and the third residue of an acceptor stem (4-3 cross-links). Some bacterial peptidoglycans also contain 3-3 cross-links that are formed by another class of enzymes called L,D-transpeptidases. In this work, we investigate the formation of unusual bacterial 1-3 peptidoglycan cross-links. We describe a version of the PGFinder software which can identify 1-3 cross-links and report the high-resolution peptidoglycan structure of *Gluconobacter oxydans* (a model organism within the *Acetobacteraceae* family). We reveal that *G. oxydans* peptidoglycan contains peptide stems made of a single alanine as well as several dipeptide stems with unusual amino acids at their C-terminus. Using a Sudoku transposon library, we identified a *G. oxydans* mutant with a drastic reduction in 1-3 cross-links. Through complementation experiments in *G. oxydans* and recombinant protein production in a heterologous host, we identify an L,D-transpeptidase enzyme with a domain distantly related to the YkuD domain responsible for these non-canonical reactions. This work revisits the enzymatic capabilities of L,D-transpeptidases, a versatile family of enzymes that play a key role in bacterial peptidoglycan remodelling.

## Introduction

Peptidoglycan is an essential component of the bacterial cell envelope that confers cell shape and resistance to a high internal osmotic pressure (1). This bag-shaped macromolecule surrounding the cytoplasmic membrane is made of disaccharide-peptides as building blocks. Their polymerization forms glycan chains alternating *N*-acetylglucosamine and *N*-acetylmuramic acid residues, substituted by short pentapeptide stems containing L- and D-amino acids (2). Depending on the bacterial species considered, the composition of peptidoglycan building block can vary (2), but in most bacteria (including *Escherichia coli*), pentapeptide stems are made of the sequence L-Ala-isoD-Glu-*meso*-DAP-D-Ala-D-Ala, (where DAP is diaminopimelic acid).

The polymerization of peptidoglycan has been extensively studied since the late 50’s, when it was discovered that this process is inhibited by penicillin, beta-lactam antibiotics widely used to combat infections (3,4). The ubiquitous enzymes that polymerize peptidoglycan, D,D-transpeptidases, are also called Penicillin Binding Proteins (PBPs). They recognize the C-terminal D-Ala-D-Ala extremity of a donor peptide stem, form an acyl-enzyme intermediate with the diamino acid in position 4, and then link this residue to the side-chain amino group of the amino acid in position 3 of an acceptor stem (4-3 cross-link). Beta-lactams are structural analogues of the D-Ala-D-Ala stems and can be used as suicide substrates (5), leading to growth arrest and cell death (6). Alternative 3-3 peptidoglycan cross-links were originally described in *Mycobacteria* (7). These types of bonds are prevalent in the peptidoglycan of important pathogens such as *Mycobacterium tuberculosis* (8), *Mycobacterium leprae* (9) and *Clostridium difficile* (10). In *Enterococcus faecium*, resistance to beta-lactams and glycopeptides can emerge when 4-3 cross-links are replaced by 3-3 cross-links. The complete bypass of the D,D-transpeptidation pathway in *E. faecium* led to the identification of the enzyme catalysing the formation of 3-3 bonds (11) which is an L,D-transpeptidase. Instead of recognizing the D-Ala-D-Ala extremity of the pentapeptide donor stem, L,D-transpeptidases use a tetrapeptide stem as a substrate. These enzymes can perform several activities depending on the substrate they use as an acceptor and can act as a carboxypeptidase (cleaving the fourth residue of the donor stem) (12), a transpeptidase (forming 3-3 cross-linked muropeptides or covalently anchoring proteins to peptidoglycan) (13,14), or an endopeptidase (cleaving 3-3 cross-links or the link between peptidoglycan and covalently attached proteins) (15–17). Finally, L,D-transpeptidases can also exchange the fourth amino-acid of a peptide stem for another amino-acid (18,19). The peptidoglycan structural changes catalysed by L,D-transpeptidases (called remodelling) plays an important role in cell shape (12), resistance to abiotic stress (20), pathogenesis, and host immunity (21).

A recent study described the existence of peptidoglycan 1-3 cross-links in *Acetobacteraceae* and proposed that this unusual type of cross-link could play a role for the survival of these organisms in the context of their interaction with the fly immune system and during competition with other organisms (22). In this work, we describe a version of PGFinder that can automate the analysis of peptidoglycans with 1-3 cross-links. Using this tool, we determine the high-resolution structure of *Gluconobacter oxydans* peptidoglycan and reveal that it contains a high proportion of previously undescribed disaccharide-dipeptides with non-canonical amino acids at their C-terminus. Using a transposon mutant and its complemented derivative, as well as heterologous expression experiments, we demonstrate that *G. oxydans* 1-3 cross-links are formed by an enzyme with a domain distantly related to the YkuD domain of canonical L,D-transpeptidases. Collectively, our data show that L,D-transpeptidases have evolved to carry out enzymatic reactions using either tetrapeptide or dipeptide stems as donors.

## Results

### Building a software tool for the structural analysis of 1-3 cross-linked peptidoglycans

Prior to this study, the PGFinder software (v1.0.3; https://mesnage-org.github.io/pgfinder/) could only generate dimers and trimers crosslinked via 3-3 and 4-3 bonds (23). To perform the structural analysis of *G. oxydans* peptidoglycan, we modified PGFinder and its graphical user interface to enable the creation of dynamic databases containing dimers and trimers with 1-3 cross-links (Fig. 1). This PGFinder upgrade (v1.1.0) was tested using datasets from *G. oxydans*.

**Figure 1.**
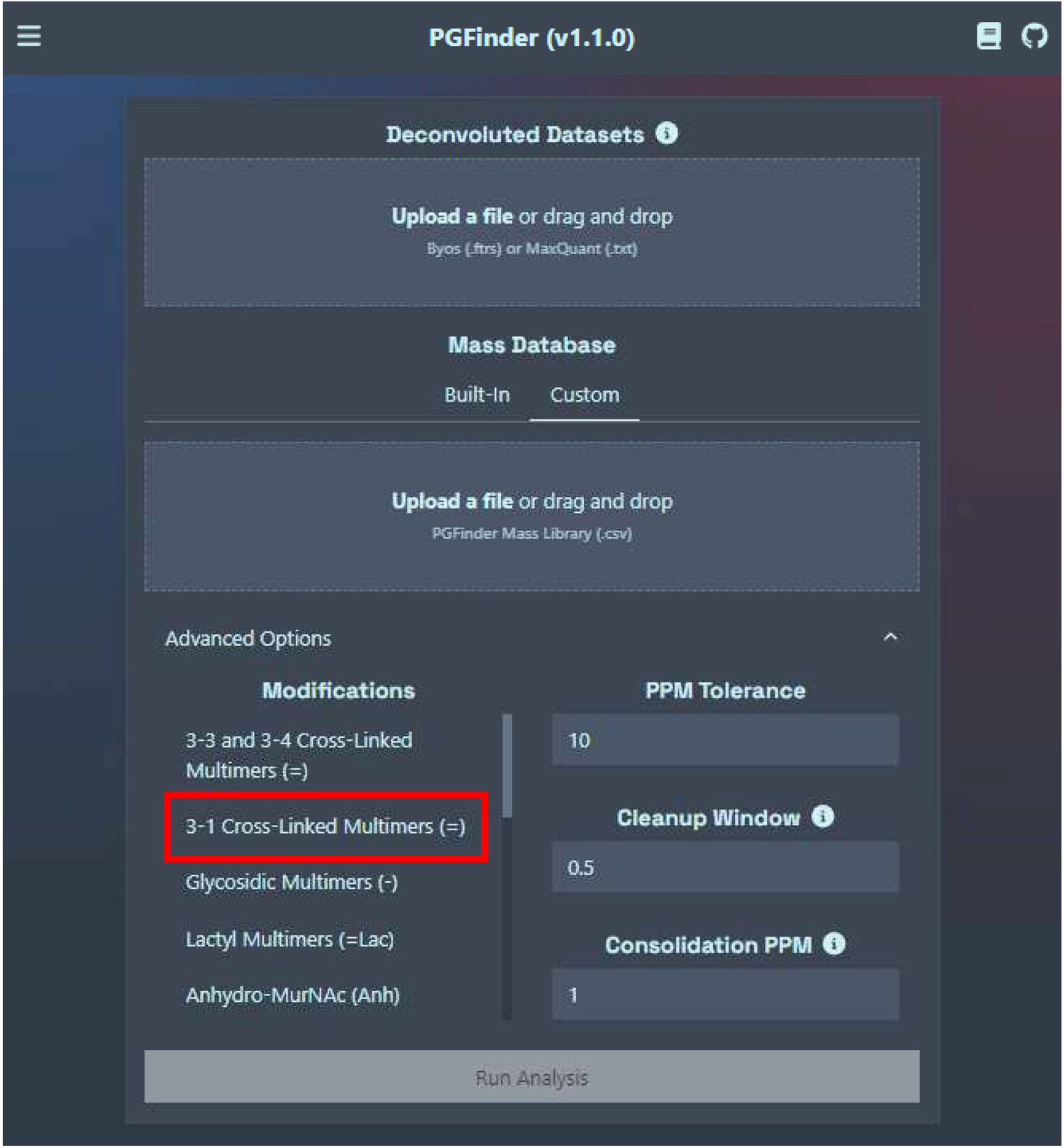
Analysis of 1-3 cross-linked peptidoglycans using PGFinder v1.1.0. The advanced option command enables the creation of dynamic databases containing 1-3 cross-linked dimers and trimers.

### High resolution analysis of *G. oxydans* B58 peptidoglycan

Peptidoglycan was purified from *G. oxydans* B58 cells harvested during both exponential (Fig. 2A) and stationary phase (Fig. 2B). As expected, the muropeptide profiles revealed changes indicative of major peptidoglycan remodelling during stationary phase. We used a combination of automated tools previously described to determine the high-resolution structure of *G. oxydans* peptidoglycan (18,23). A two-step custom search strategy was followed (Fig. S1). We first used the proprietary Byonic™ software to identify monomers based on tandem mass spectrometry data. The search space contained mono-, di-, tri-, tetra- and pentapeptides containing Alanine in position 1, glutamic acid (E) or glutamine (Q) in position 2, *meso*-diaminopimelic acid (J) or amidated *meso*-diaminopimelic acid (Z) in position 3, any possible amino acids (X) in position 4, and pentapeptides containing AX dipeptides at their C-terminus (Table S1).

**Figure 2.**
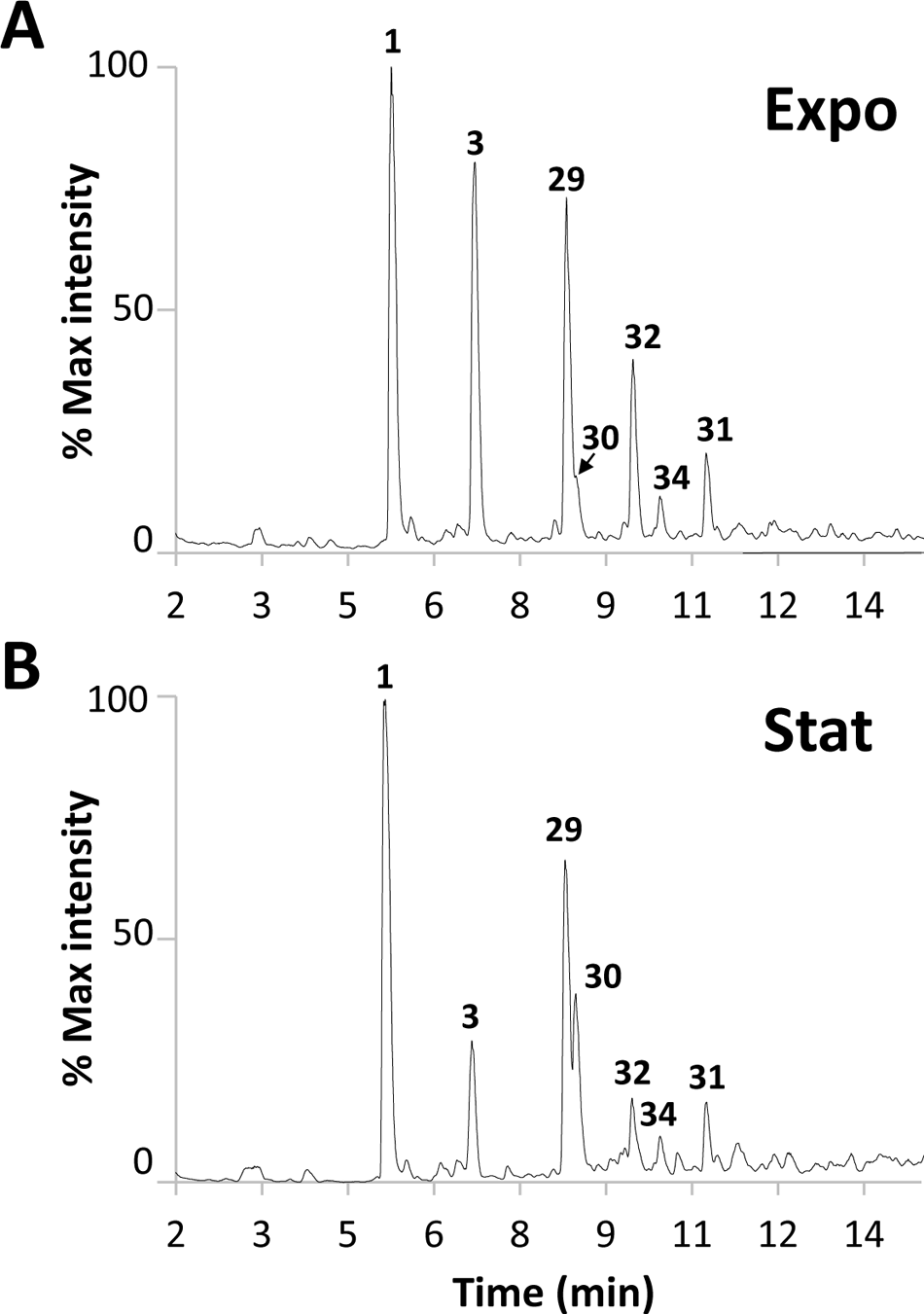
HPLC-MS chromatogram of *G. oxydans* reduced disaccharide-peptides. Strain B58 was grown in YPM media to exponential (**A**) or stationary phase (**B**). The numbers refer to the muropeptide structures described in Table 1.

**Table 1.** *G. oxydans* peptidoglycan composition.

|  | Muropeptide <sup>a</sup> | WT |  | Ldt<sub>Go1</sub> | Ldt<sub>Go2</sub> | RT (min)<sup>b</sup> | Theoretical mass (Da) | Δppm |
| --- | --- | --- | --- | --- | --- | --- | --- | --- |
|  |  | Expo | Stat |  |  |  |  |  |
| 1 | gm-AEJ<sub>NH2</sub> | 36.519% | 41.440% | 48.001% | 34.926% | 5.32 ± 0.05 | 869.3867 | 4.01 |
| 2 | gm-AE | 4.239% | 10.518% | 9.628% | 6.887% | 6.79 ± 0.03 | 698.2859 | 2.77 |
| 3 | gm-AEJA | 22.572% | 3.270% | 4.019% | 28.131% | 6.80 ± 0.01 | 941.4078 | 3.38 |
| 4 | gm-AE (Anh) | 0.880% | 1.158% | 0.656% | 0.505% | 11.06 ± 0.00 | 678.2597 | 1.52 |
| 5 | gm-AEJ<sub>NH2</sub> (Anh) | 0.522% | 0.682% | 0.472% | 0.161% | 8.61 ± 0.01 | 849.3605 | 2.29 |
| 6 | gm-AEJ<sub>NH2</sub>A | 0.254% | 0.290% | 0.343% | 0.012% | 6.37 ± 0.27 | 940.4238 | 1.39 |
| 7 | gm-AEJ | 0.711% | 0.356% | 0.193% | 0.014% | 5.66 ± 0.04 | 870.3707 | 2.14 |
| 8 | gm-AEJAG | 0.494% | 0.064% | 0.130% | 0.247% | 6.49 ± 0.02 | 998.4293 | 1.46 |
| 9 | gm-AF | 0.072% | 0.171% | 0.096% | ND | 14.54 ± 0.01 | 716.3117 | 0.89 |
| 10 | gm-AEJAA | 0.353% | 0.074% | 0.094% | 0.067% | 8.08 ± 0.39 | 1012.4450 | 2.97 |
| 11 | gm-A | 0.335% | 1.385% | 0.534% | 0.024% | 6.26 ± 0.03 | 569.2433 | 1.8 |
| 12 | gm-AEJA (Anh) | 0.425% | 0.093% | 0.079% | 0.131% | 10.42 ± 0.01 | 921.3816 | 1.45 |
| 13 | gm-AEJAK | 0.114% | 0.021% | 0.042% | 0.057% | 5.86 ± 0.03 | 1069.5028 | 1.07 |
| 14 | gm-AY | 0.032% | 0.100% | 0.042% | ND | 10.72 ± 0.01 | 732.3066 | 0.52 |
| 15 | gm-AI | 0.027% | 0.070% | 0.038% | ND | 12.89 ± 0.00 | 682.3274 | 0.81 |
| 16 | gm-AI (Anh) | ND | 0.029% | ND | ND | 12.75 | 662.3012 | 0.99 |
| 17 | gm-AEJAR | 0.121% | 0.028% | 0.037% | 0.062% | 6.42 ± 0.03 | 1097.5089 | 1.57 |
| 18 | gm-AEJAH | 0.100% | 0.035% | 0.035% | 0.050% | 5.99 ± 0.03 | 1078.4667 | 1.41 |
| 19 | gm-AEJAN | 0.022% | 0.069% | 0.033% | 0.038% | 10.37 ± 4.33 | 1055.4508 | 1.72 |
| 20 | gm-AI (-Ac) | 0.055% | ND | ND | ND | 6.29 | 640.3169 | 0.24 |
| 21 | gm-AEJ<sub>NH2</sub> (-Ac) | 0.036% | 0.015% | 0.023% | 0.022% | 5.05 ± 0.06 | 827.3762 | 1.15 |
| 22 | gm-AQ | 0.057% | 0.026% | 0.022% | 0.047% | 5.77 ± 0.04 | 697.3019 | 0.97 |
| 23 | gm-AEJAE | 0.029% | 0.019% | 0.018% | 0.031% | 7.29 ± 0.01 | 1070.4504 | 0.50 |
| 24 | gm-AEJ (Anh) | 0.011% | ND | 0.017% | ND | 9.03 ± 0.00 | 850.3445 | 1.15 |
| 25 | gm-AEJA (-Ac) | 0.021% | 0.002% | 0.007% | 0.027% | 6.43 ± 0.02 | 899.3973 | 1.11 |
| 26 | gm-AEJ<sub>NH2</sub>A (Anh) | ND | ND | 0.006% | ND | 9.93 | 920.3976 | 1.17 |
| 27 | gm-AE (-Ac) | ND | 0.030% | 0.003% | ND | **5.62 ± 0.01** | 656.2754 | 0.14 |
| 28 | gm-AF (-Ac) | ND | ND | 0.003% | ND | 16.28 | 674.3012 | 1.24 |
| 29 | gm-AEJ<sub>NH2</sub>=gm-AEJA | 13.782% | 15.265% | 18.938% | 15.714% | 8.43 ± 0.00 | 1792.7836 | 3.35 |
| 30 | gm-AEJ<sub>NH2</sub>=gm-A | 2.592% | 13.391% | 7.317% | 0.037% | 8.60 ± 0.00 | 1420.6194 | 1.75 |
| 31 | gm-AEJ<sub>NH2</sub>=gm-AEJA (Anh) | 3.741% | 4.098% | 4.220% | 4.109% | 10.87 ± 0.00 | 1772.7574 | 2.38 |
| 32 | gm-AEJA=gm-AEJA | 6.802% | 2.822% | 2.318% | 6.976% | 9.59 ± 0.00 | 1864.8047 | 2.51 |
| 33 | gm-AEJ<sub>NH2</sub>=gm-A (Anh) | 0.243% | 0.794% | 0.609% | ND | 11.43 ± 0.01 | 1400.5932 | 0.85 |
| 34 | gm-AEJA=gm-A | 2.520% | 2.028% | 0.512% | 0.011% | 10.06 ± 0.00 | 1492.6405 | 1.02 |
| 35 | gm-AEJA=gm-AEJA (Anh) | 0.729% | 0.388% | 0.199% | 0.601% | 12.01 ± 0.00 | 1844.7785 | 1.37 |
| 36 | gm-AEJ=gm-AEJA | 0.186% | 0.136% | 0.127% | ND | 9.03 ± 0.05 | 1793.7676 | 0.68 |
| 37 | gm-AEJA=gm-A (Anh) | 0.317% | 0.197% | 0.074% | ND | 12.87 ± 0.02 | 1472.6143 | 0.41 |
| 38 | gm-AEJAG=gm-AEJA | 0.134% | 0.055% | 0.049% | 0.078% | 9.18 ± 0.00 | 1921.8262 | 0.54 |
| 39 | gm-AEJ=gm-AEJA (Anh) | 0.029% | 0.023% | 0.034% | ND | 11.46 ± 0.02 | 1773.7414 | 1.45 |
| 40 | gm-AEJAG=gm-A | 0.068% | 0.030% | 0.022% | 0.013% | 10.42 ± 1.48 | 1549.6617 | 3.24 |
| 41 | gm-AEJ=gm-A | 0.048% | 0.145% | 0.019% | ND | 9.38 ± 0.00 | 1421.6034 | 0.94 |
| 42 | gm-AEJA=gm-AEJA (Anh) | ND | 0.014% | ND | ND | 9.84 | 1844.7786 | 0.09 |
| 43 | gm-AEJAA=gm-AEJA | 0.034% | 0.009% | 0.016% | 0.023% | 9.84 ± 0.01 | 1935.8419 | 0.67 |
| 44 | gm-AEJ<sub>NH2</sub>A=gm-AEJA | 0.007% | 0.008% | 0.012% | ND | 9.32 ± 0.04 | 1863.8207 | 0.70 |
| 45 | gm-AEJAG=gm-AEJA (Anh) | 0.032% | 0.014% | 0.010% | 0.016% | 11.53 ± 0.01 | 1901.8000 | 1.14 |
| 46 | gm-AEJAR=gm-AEJA | 0.017% | ND | 0.006% | 0.007% | **9.00 ± 0.00** | 2020.9058 | 0.32 |
| 47 | gm-AEJAN=gm-AEJA | ND | ND | 0.005% | 0.004% | **8.76 ± 0.01** | 1978.8477 | 0.81 |
| 48 | gm-AEJ<sub>NH2</sub>A=gm-AEJA (Anh) | ND | ND | 0.004% | ND | 11.56 | 1843.7945 | 0.37 |
| 49 | gm-AEJ<sub>NH2</sub>=gm-AEJA (-Ac) | 0.005% | ND | 0.003% | 0.004% | 8.18 ± 0.03 | 1750.7731 | 2.03 |
| 50 | gm-AEJAH=gm-AEJA | 0.013% | ND | ND | 0.006% | **8.64 ± 0.00** | 2001.8636 | 0.74 |
| 51 | gm-AEJ<sub>NH2</sub>A=gm-A |  | ND | ND | 0.005% | 16.44 | 1491.6565 | 0.55 |
| 52 | gm-AEJAA=gm-AEJA (Anh) | 0.006% | ND | 0.002% | 0.004% | 12.12 ± 0.01 | 1915.8157 | 0.63 |
| 53 | gm-AEJAG=gm-A (Anh) | 0.005% | ND | 0.002% | ND | **9.71 ± 2.72** | 1529.6355 | 1.74 |
| 54 | gm-AEJAR=gm-AEJA (Anh) | 0.004% | ND | ND | ND | 11.24 | 2000.8796 | 0.61 |
| 55 | gm-AEJ<sub>NH2</sub>=gm-AEJA=gm-AEJA | 0.370% | 0.384% | 0.551% | 0.554% | 9.97 ± 0.00 | 2716.1805 | 0.96 |
| 56 | gm-AEJ<sub>NH2</sub>=gm-AEJA=gm-AEJA (Anh) | 0.180% | 0.196% | 0.210% | 0.229% | 11.84 ± 0.00 | 2696.1543 | 0.78 |
| 57 | gm-AEJA=gm-AEJA=gm-AEJA | 0.099% | 0.045% | 0.119% | 0.136% | 10.84 ± 0.00 | 2788.2016 | 1.89 |
| 58 | gm-AEJA=gm-AEJA=gm-AEJA (Anh) | 0.032% | 0.015% | 0.024% | 0.033% | 12.67 ± 0.03 | 2768.1754 | 0.28 |
| 59 | gm-AEJ=gm-AEJA=gm-AEJA | ND | ND | 0.019% | ND | **10.59 ± 0.00** | 2717.1645 | 1.18 |
| 60 | gm-AEJ=gm-AEJA=gm-AEJA (Anh) | ND | ND | 0.005% | ND | 12.51 | 2697.1383 | 0.52 |
| 61 | gm-AEJAG=gm-AEJA=gm-AEJA | 0.005% | ND | 0.004% | 0.003% | 10.49 ± 0.01 | 2845.2231 | 0.53 |

<sup>a</sup> g, GlcNAc; m, MurNAc; A, Alanine; E, isoglutamic acid; J, *meso*-diaminopimelic acid; J<sub>NH2</sub>, amidated *meso*-diaminopimelic acid
<sup>b</sup> standard deviations in bold are determined from 2 values only

Seventeen monomers showing more than half of the expected *b* and *y* ions in their fragmentation spectra were identified by Byonic^TM^ (Fig. S2). Interestingly, these included several muropeptides with a dipeptide stem other than AE that were not previously identified and a lack of tetrapeptide stems with unusual amino acids formed by canonical L,D-transpeptidases. In addition to amidated *meso*-DAP residues, we also found the presence of deacetylated GlcNAc residues (glucosamine) which were not previously reported. The disaccharide-peptides corresponding to these validated monomers were combined to create the database called DB0_Go (Table S2). We next performed a PGFinder search, enabling the identification of dimers and trimers with 4-3, 3-3, and 1-3 cross-links as well as modified disaccharides (deacetylated and containing MurNAc residues). A total of 61 masses matching the monoisotopic mass of theoretical structures were identified (Table 1), revealing a far more complex structure than previously reported. A direct comparison of the peptidoglycan from cells harvested during exponential and stationary phase showed an increased cross-linking index (19.9 % versus 15.9 %), partly due to an increased proportion of 1-3 cross-links in the stationary phase (16.6% versus 5.8 %). The higher proportion of 1-3 cross-links was concomitant with the higher proportion of disaccharide-dipeptide structures detected in stationary phase (13.5% versus 5.7%). Very little variation was observed in the glycan chain length between exponential and stationary phase, with the average length being equal to 24 and 22 disaccharides, respectively.

By analogy with the transpeptidation reaction leading to the formation of 3-3 bonds (Fig. 3A), we hypothesized that the formation of 1-3 bonds uses muropeptides with a dipeptide stem as donor substrates (Fig. 3B). According to this hypothesis, the enzyme is predicted to form an acyl enzyme intermediate with a disaccharide-alanine.

**Figure 3.**
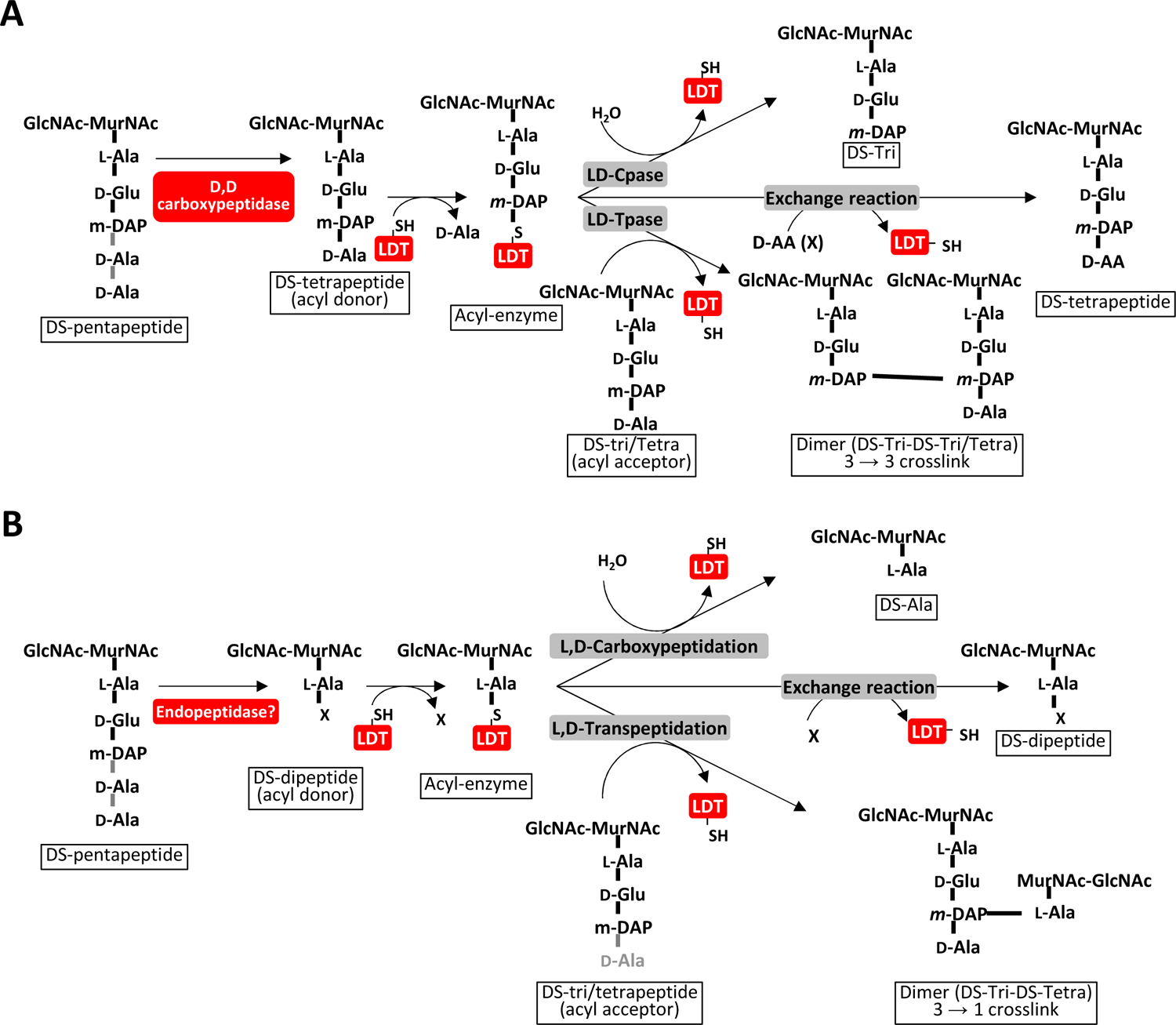
Schematic representation of the L,D-transpeptidation reactions leading to the formation of 3-3 and 1-3 cross-links. The enzymatic reactions carried out by L,D-transpeptidases in organisms with 3-3 cross-links are described (**A**). By analogy with these L,D transpeptidation reactions, we propose a model that leads to distinct reactions in *G. oxydans* (**B**). We hypothesize that an unidentified endopeptidase generates disaccharide dipeptides. These muropeptides are used as substrates to form an acyl-enzyme intermediate. Depending on the acceptor group, the reaction can lead to a carboxypeptidase reaction or a transpeptidation reaction that generates either a dimer or a disaccharide-dipeptide. GlcNAc, *N*-acetylglucosamine; MurNAc, *N*-acetylmuramic acid; LDT, L,D-transpeptidase; DS, disaccharide (GlcNAc-MurNAc); X, any D amino acid; DS-Tri, disaccharide-tripeptide; DS-Tetra, disaccharide-tetrapeptide.

### Identification of the L,D-transpeptidase catalysing the formation of 1-3 cross-links in *G. oxydans*

Interestingly, *G. oxydans* peptidoglycan does not contain any tetrapeptide stems with unusual amino acids at their C-terminus, which are characteristic of canonical L,D-transpeptidase enzymatic activity (Fig. 3A). We therefore hypothesized that in *G. oxydans*, L,D-transpeptidases could perform a similar enzymatic reaction using disaccharide-dipeptides as substrates instead (Fig. 3B). We searched the *G. oxydans* genome to identify genes encoding homologs of the L,D-transpeptidases and found two putative L,D-transpeptidases (labeled GOX1074, 337 residues and GOX2269, 171 residues in *G. oxydans* 621H) related to the YkuD catalytic domain (Pfam: PF03734). We hypothesized that one or both enzymes could catalyze the formation of 1-3 cross-links and renamed these putative L,D-transpeptidases Ldt_Go1_ and Ldt_Go2_. To test this, we took advantage of the *G. oxydans* B58 Sudoku library previously described (24) and analyzed the peptidoglycan structure of the two transposon mutants with an insertion in each of the *ldt_Go_* genes by LC-MS. Comparison of the TIC profiles indicated the presence of a peak corresponding to the major 1-3 dimer (gm-AEJ_NH2_=gm-A) in the wild type (Fig. 4A) and *ldt_Go1_* mutant (Fig. 4B), whilst no equivalent peak was detected in the *ldt_Go2_* mutant (Fig. 4C). Analysis of the extracted ion chromatograms for all molecules eluted between 7.5 and 11.5 min revealed a drastic reduction of 1-3 cross-links in the *ldt_Go2_* peptidoglycan sample (0.035% versus 7.3% in the *ldt_Go1_*mutant). The *ldt_Go2_* mutation was also associated with a reduction of disaccharide-dipeptides and 1-3 crosslinked dimers as compared to the parental strain and the *ldt_Go1_* mutant (Table 1). Collectively, our LC-MS data showed that Ldt_Go2_ plays a major role in the unusual L,D-transpeptidation reactions in *G. oxydans*, including the formation of 1-3 cross-links.

**Figure 4.**
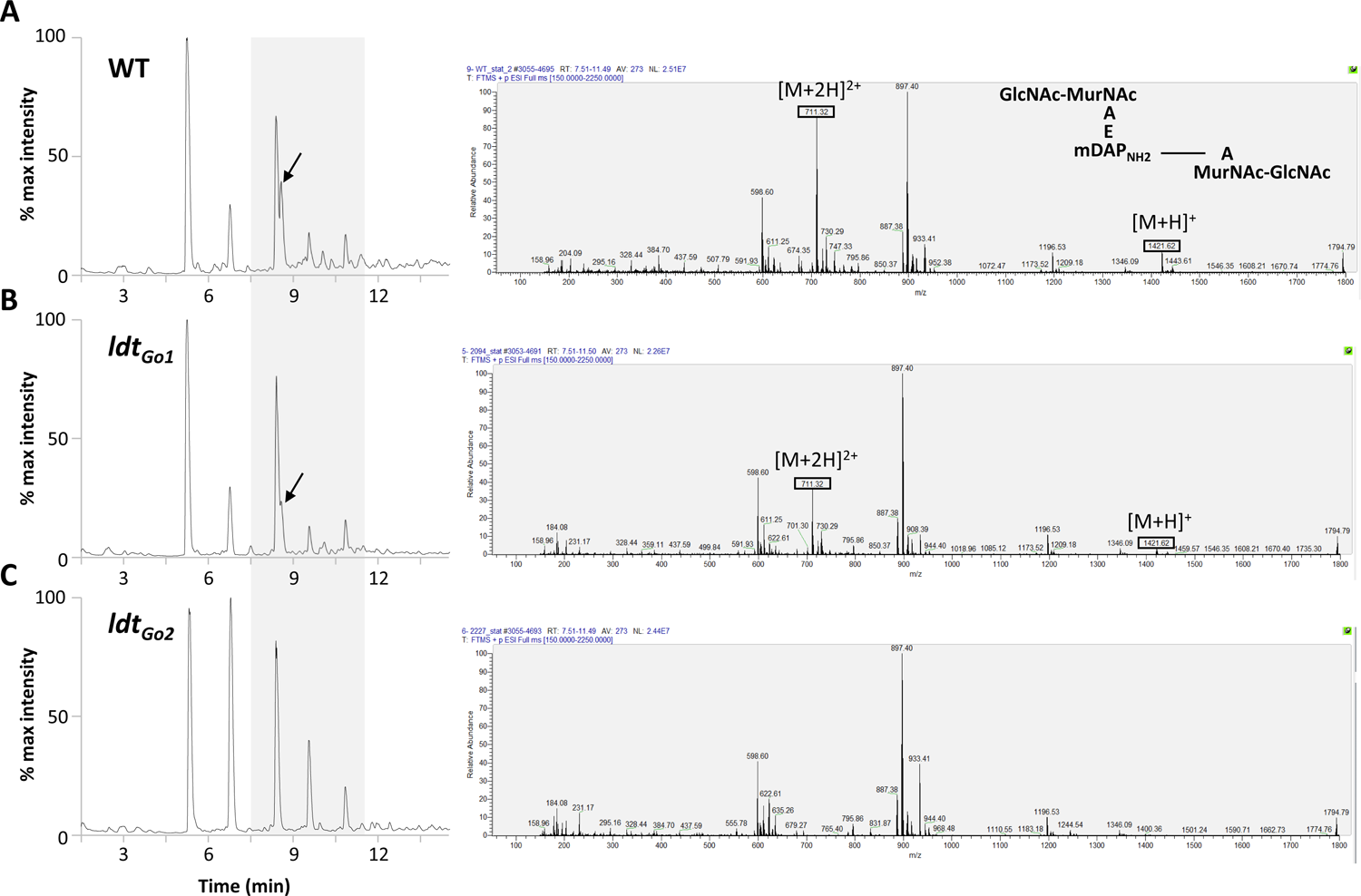
LC-MS detection of 2-1 cross-links in *G. oxydans* peptidoglycan. TIC of *G. oxydans* B58 (**A**) and mutants with a transposon insertion in the *ldt_Go1_* (**B**) and *ldt_Go2_* genes (**C**) are shown on the left hand side. Extracted ions corresponding to the muropeptides eluted between 7.5 and 11.5 min are shown on the right hand side. The major dimer with a 1-3 cross-link (shown with an arrow on the TIC and on the top MS spectrum) is associated with two major protonated ions: a singly charged ion with an *m/z* at 1421.32 and a doubly charged ion with an *m/z* at 711.32. None of these ions were detected in the peptidoglycan from the *ldt_Go2_* mutant, demonstrating that this gene is essential for the formation of 1-3 cross-links.

### Complementation and heterologous expression experiments show that the Ldt_Go2_ enzyme is sufficient to catalyse peptidoglycan 1-3 cross-links

To verify that the drastic reduction of 1-3 cross-links was associated with the disruption of *ldt_Go2_* and not a secondary mutation, we built a complementation strain expressing Ldt_Go2_ under the anhydrotetracycline-inducible promoter (25). The production of Ldt_Go2_ in the *ldt_Go2_*transposon mutant background clearly restored the presence of a peak corresponding to the major 1-3 cross-linked dimers (Fig. S3).

We further confirmed the enzymatic activity of Ldt_Go2_ by producing the full-length protein in *E. coli*. Since *E. coli* peptidoglycan contains disaccharide-dipeptides (gm-AE) that represent the substrate for the 1-3 transpeptidation reaction, we anticipated that recombinant Ldt_Go2_ could generate the expected products found in *G. oxydans* in this heterologous host. Peptidoglycan was purified from *E. coli* transformed with either the empty pET expression vector or a recombinant derivative expressing *ldt_Go2_*, digested with mutanolysin, and analysed by reversed phase HPLC (Fig. 5A, top and bottom trace, respectively). A simple search strategy was followed to identify and quantify muropeptides resulting from unusual L,D-transpeptidation reactions (Fig. S4).

**Figure 5.**
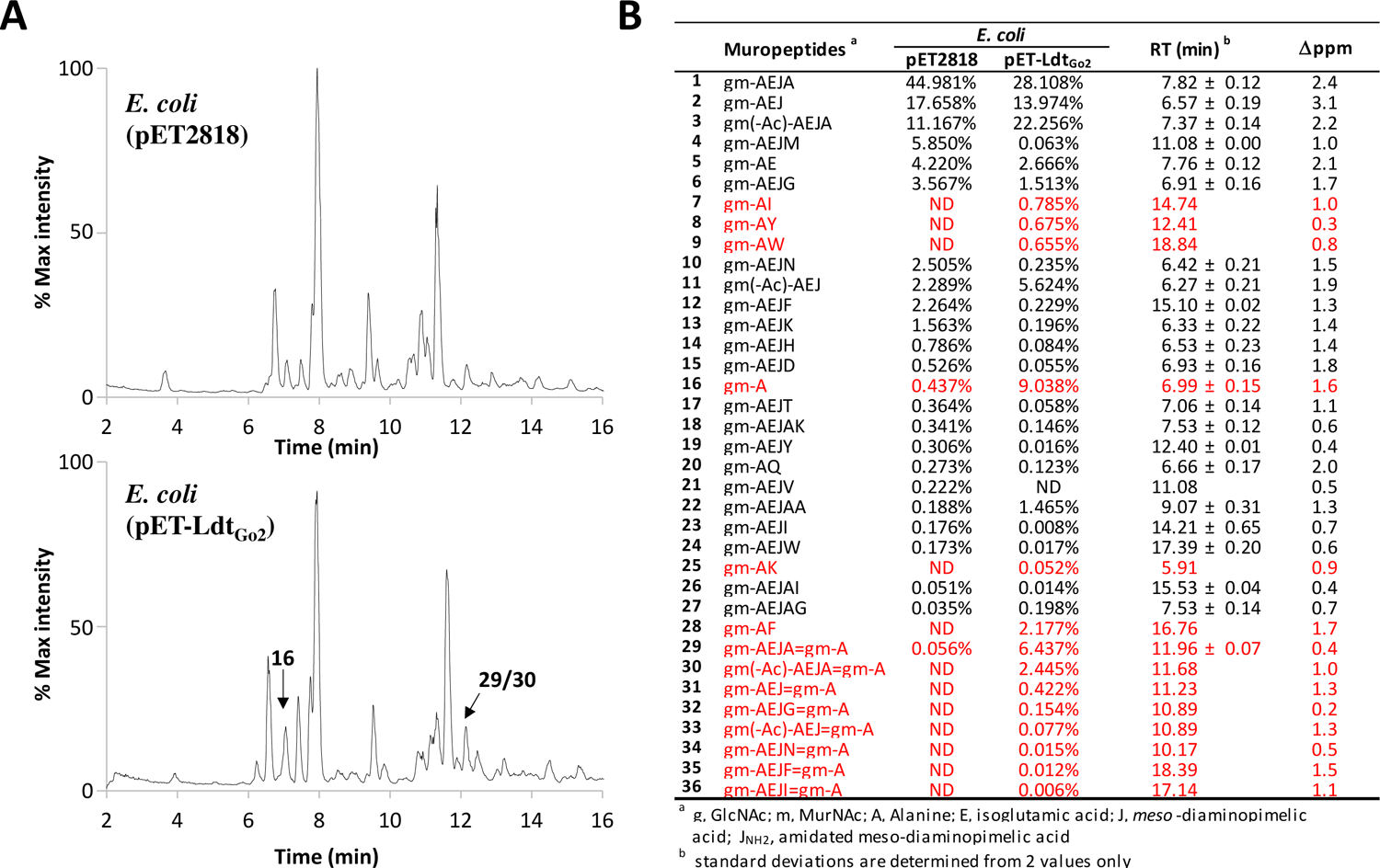
Heterologous protein synthesis of Ldt_Go2_ in *E. coli* BL21(DE3) increases the proportion of 1-3 L,D-transpeptidation products. *E. coli* BL21(DE3) transformed with the control pET2818 plasmid or pET2818 encoding Ldt_Go2_ was grown in auto-induction medium overnight and peptidoglycan from both cultures were purified. The muropeptide profile are shown in (**A**); bottom profile is from the control strain, bottom panel is from *E. coli* expressing recombinant Ldt_Go2_. Two major peaks containing muropeptides of interest resulting from 1-3 L,D-transpeptidation are indicated. (**B**) PGFinder analysis of *E. coli* control strain (transformed with the empty plasmid, *E. coli* (pET2818)) and expressing Ldt_Go2_ (*E. coli* (pET-Ldt_Go2_)). Only monomers validated by Byonic based on MS/MS data were search as well as their 1-3 transpeptidation products. The monomers and dimers resulting from 1-3 L,D-transpeptidation are indicated in red.

First, we identified monomers based on MS/MS data using the Byonic™ module from Byos®. We searched for all possible disaccharide-peptides containing one to five amino acids (A, AX, AEJ, AEJX and AEJAX, where J is *meso*-diaminopimelic acid and X any amino acid) adding sugar deacetylation previously identified in *E. coli* (23) as a potential glycan modification (Table S3). Twenty-eight monomers validated by MS/MS analysis were selected to create a database called DB0_Ec (Table S4). This monomer database was then run through PGFinder to identify, compare, and quantify muropeptides in the *E. coli* expression strain and its derivative expressing Ldt_Go2_. To focus on 1-3 L,D transpeptidation products, we only enabled the search for 1-3 dimers which contain the gm-A moiety. Interestingly, the PGFinder search revealed a very low amount of 1-3 transpeptidation products in *E. coli* (gm-A, 0.44% and gm-AEJA=gm-A, 0.056%) (Fig. 5B). A striking increase in gm-A (9.0%), gm-AX (4.3%) monomers and dimers resulting from 1-3 cross-linking (9.5%) was detected in the peptidoglycan of the strain expressing Ldt_Go2_. demonstrating that this enzyme is an L,D-transpeptidase that can perform all the reactions described in Fig. 3 (carboxypeptidation, exchange and 1-3 transpeptidation).

### Ldt_Go2_ is characterised by an atypical YkuD-like catalytic domain that can be found in distant families of bacteria with 1-3 peptidoglycan cross-links

To place *G. oxydans*’s L,D-transpeptidases in a broader evolutionary context, homologues were extracted from genomes of numerous alphaproteobacterial species (Table S5), including those previously shown to contain 1-3 peptidoglycan cross-links (22). Four other organisms with characterized L,D-transpeptidases (*E. coli*, *C. difficile, M. tuberculosis* and *E. faecium*) were added. Phylogenetic reconstruction of all putative alphaproteobacterial L,D-transpeptidases revealed that both Ldt_Go1_ and Ldt_Go2_ homologues form distinct clades representing previously uncharacterised transpeptidase subfamilies (Fig. 6A).

**Figure 6.**
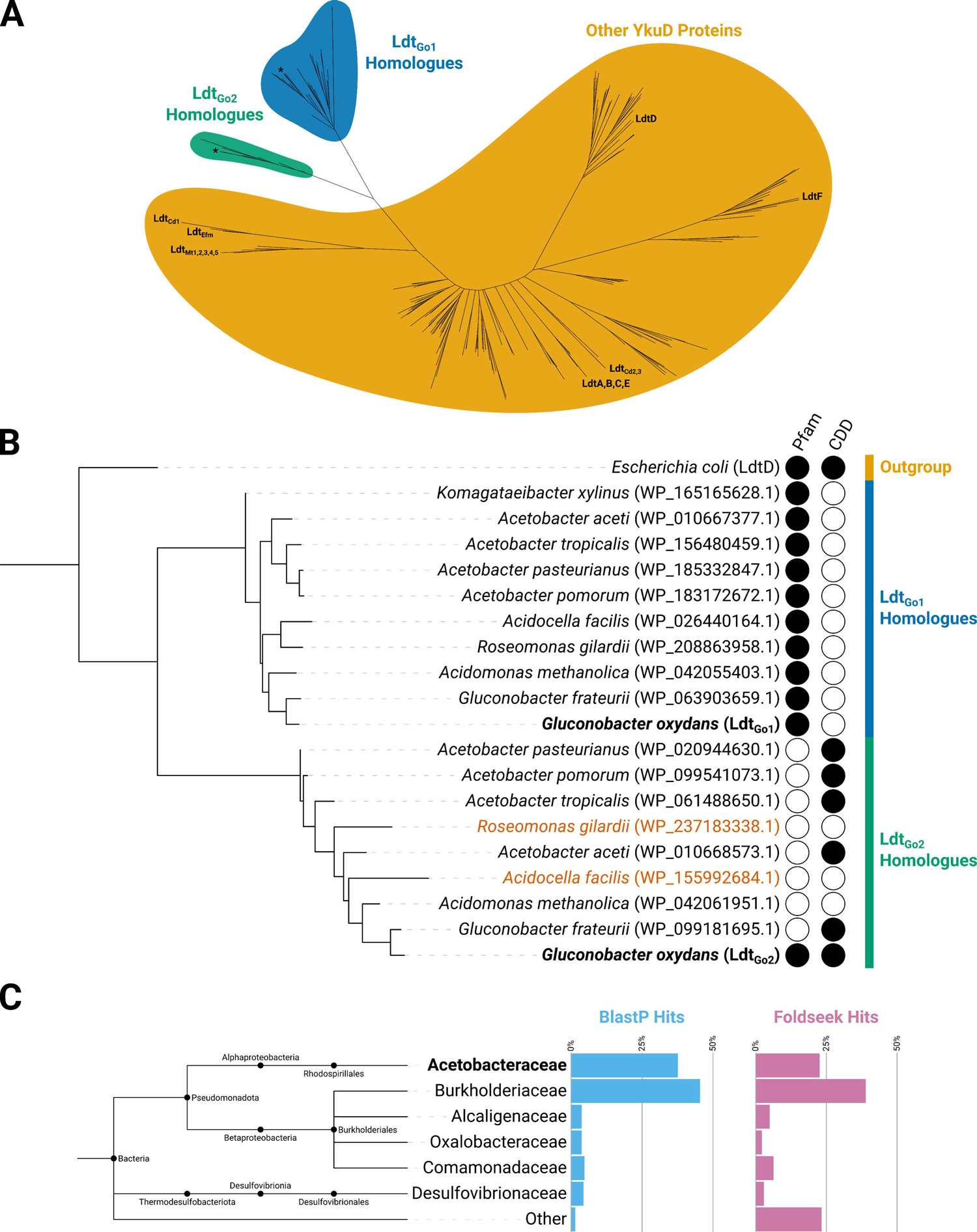
Ldt_Go2_ represents a distinct L,D-transpeptidase subclade with a divergent catalytic domain that is found primarily in the *Acetobacteraceae* and *Burkholderiaceae*. (**A**) An unrooted phylogenetic tree of putative L,D-transpeptidases throughout Alphaproteobacteria and characterized enzymes (Table S5) reveals that Ldt_Go1_ and Ldt_Go2_ homologues form distinct transpeptidase subfamilies. Ldt_Go1_ and Ldt_Go2_ are labelled with asterisks (*). Previously characterised L,D-transpeptidases from *Escherichia coli* (LdtA-F), *Clostridioides difficile* (Ldt_Cd1-3_), *Mycobacterium tuberculosis* (Ldt_Mt1-5_) and *Enterococcus faecium* (Ldt_Efm_) have also been labelled. (**B**) A phylogram of Ldt_Go1_, Ldt_Go2_, and their homologues reveals that whilst all Ldt_Go1_ homologues were annotated with the canonical L,D-transpeptidase Pfam domain (YkuD) (42), most Ldt_Go2_ homologues were annotated only with the CDD YkuD_like domain (43), and the rest lacked domain annotation entirely. The Ldt_Go2_ homologues highlighted in orange are found in bacterial species where no 1-3 cross-links could be detected (22). (**C**) An expanded search for Ldt_Go2_ homologues beyond the Alphaproteobacteria reveals the presence of this subfamily throughout the Burkholderiales and *Desulfovibrionaceae*. Structural homologues located using Foldseek (28) show a similar evolutionary distribution as those located using BlastP (26).

When annotated using InterProScan and its default significance thresholds, Ldt_Go1_ homologues are shown to contain a canonical YkuD (Pfam: PF03734) domain, but the more distantly related Ldt_Go2_ homologues typically lack this annotation. Instead, most are annotated with a YkuD-like (CDD: cd16913) domain, and others contain no domain annotations at all (Fig. 6B). Although Ldt_Go2_ is annotated with a canonical YkuD domain, it should be noted that the E-value for this annotation is high (2.0e-1), indicative of a significant divergence from the canonical YkuD domain.

Finally, to better understand the distribution of this unusual L,D-transpeptidase subfamily beyond the Alphaproteobacteria, the catalytic domain of Ldt_Go2_ was searched against the entirety of the NCBI RefSeq Select database (26). Out of the 307 hits returned from unique bacterial species, roughly 37% could be attributed to *Acetobacteraceae* like *G. oxydans*, but an even greater percentage of hits (45%) came from the *Burkholderiaceae* (Fig. 6C). Though the *Acetobacteraceae* and *Burkholderiaceae* encompass the majority of Ldt_Go2_ homologues, others are found sprinkled throughout the broader Burkholderiales and even beyond the Pseudomonadota, with homologues in the *Desulfovibrionaceae*. Since active site geometry is thought to be a key determinant of L,D-transpeptidase substrate preference and activity, a further search for structural homologues was conducted using Foldseek and an AlphaFold model of the Ldt_Go2_ catalytic domain (27,28). Setting an E-value threshold of <=2e-2 (selecting for matches better than *Bacillus subtilis*’ prototypical YkuD domain), led to 147 hits. The results of this structural search largely validated the results of the sequence-based BlastP search, with 22% of hits coming from the *Acetobacteraceae* and 39% from the *Burkholderiaceae*, but a much larger number of hits (23%) now fell outside of the families found by BLAST.

Overall, these analyses establish that L,D-transpeptidases associated with 1-3 cross-linking contain a catalytic domain related to the canonical YkuD transpeptidase domain, but form a distinct enzymatic subfamily.

## Discussion

In this study, we determine the high-resolution structure of *G. oxydans* peptidoglycan using a version of PGFinder that can generate dynamic databases containing 1-3 cross-linked multimers. We show that *G. oxydans* peptidoglycan contains a high proportion of dipeptide stems with unusual amino acids at their C-terminus, leading us to propose that the enzyme forming 1-3 cross-links uses dipeptide stems as a donor substrate. We identify two enzymes distantly related to L,D-transpeptidases making 3-3 cross-links. Based on the characterization of a transposon mutant and heterologous expression experiments, we demonstrate that one of these two candidates (Ldt_Go2_) catalyses the formation of 1-3 cross-links.

This work demonstrated that Ldt_Go2_ plays a predominant role in the formation of 1-3 cross-links in *G. oxydans*. The role of Ldt_Go1_ remains unclear since the inactivation of the corresponding gene is associated with only marginal changes in the peptidoglycan composition (Table 1). Our attempts to express recombinant Ldt_Go1_ and Ldt_Go2_ in *E. coli* as his-tagged or maltose-binding fusion proteins remained unsuccessful and both proteins were systematically found in the insoluble fraction, irrespective of the expression strains and conditions tested. Further experiments are therefore required to produce and purify these recombinant proteins to examine their activity in vitro more closely.

The formation of 3-3 cross-links in Enterococci is controlled by the availability of disaccharide-tetrapeptides used as donor substrate. In *E. faecium*, L,D-transpeptidation can bypass the D,D-transpeptidation following the activation of a cryptic D,D-carboxypeptidase (29). How the disaccharide-dipeptide substrates are generated in *G. oxydans* remains unknown. *G. oxydans* encodes two potential endopeptidases containing a CHAP domain that could generate Ldt_Go2_ substrates (GOX_RS06930 and GOX_RS07380 in *G. oxydans* 621H). The transposon inactivation of each gene was tested but did not abolish the production of 1-3 cross-links (data not shown), indicating that these genes do not play a predominant role in the formation of dipeptide stems or are functionally redundant. The inactivation of both genes simultaneously will be required to further investigate the Ldt_Go2_ partners that contribute to the formation of unusual cross-links.

Interestingly, only three Ldt_Go2_ homologs from *Roseomonas gilardii, Acidocella facilis, and Acidomonas methanolica* did not contain any YkuD-like (CDD: cd16913) catalytic domains. Although 1-3 cross-links have only been reported in *Acidomonas methanolica* (22), it would be worth revisiting the peptidoglycan in the two other species to confirm the absence of 1-3 cross-links using PGFinder. This work confirms previous studies which showed that this software is a powerful tool to elucidate the high-resolution of bacterial peptidoglycan structures and their quantification. Unlike peptidoglycan analyses based on UV quantification, PGFinder allows the systematic and unbiased identification of low abundance muropeptides accounting for less than 0.01% of all muropeptides. A striking result illustrating the low detection threshold provided by PGFinder is the identification of 1-3 crosslinks and gm-A muropeptides in *E. coli*, indicating that this organisms’ L,D-transpeptidases can also form unusual reactions.

Moving from sequence to structural analysis, the predicted fold of Ldt_Go2_ revealed the presence of a much more open, bowl-like active site (Fig. 7). Given that this enzyme uses a shorter peptide stem as a donor substrate, it is likely that the catalytic site does not require the canonical cleft or trapping loops to accommodate the substrate. Instead, the open conformation of the catalytic site could ensure that the bulky sugar moieties of a dipeptide substrate don’t limit access to the catalytic cysteine residue responsible for the formation of 1-3 cross-links.

**Figure 7.**
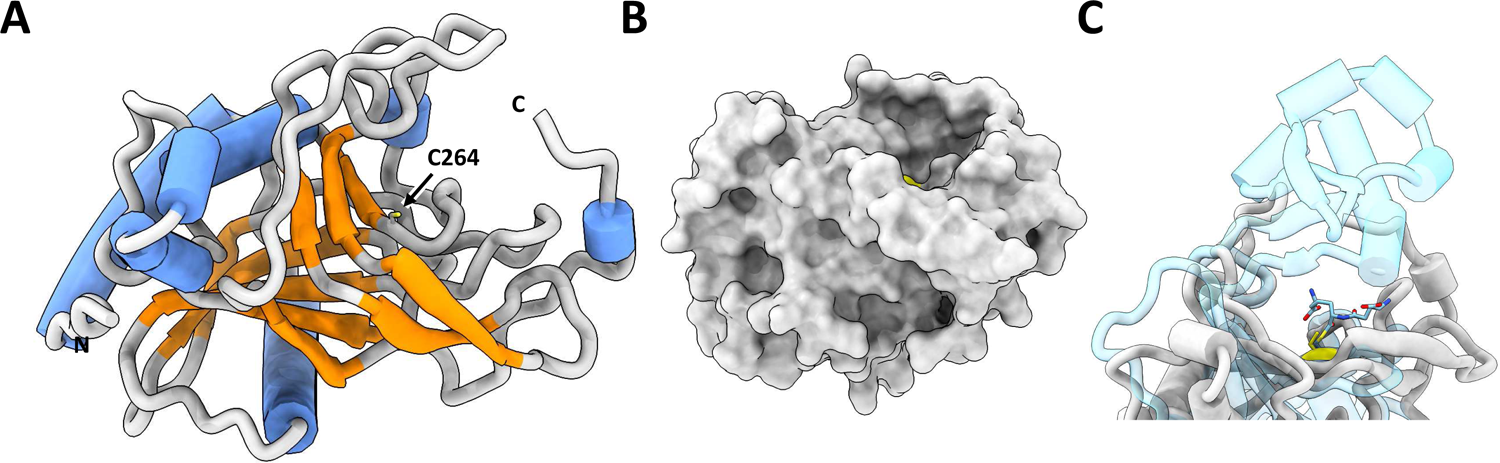
Structural analysis of G. oxydans Ldt_Go2_. (**A**) Predicted fold of Ldt_Go2_, inclusive of well-modelled residues 79 to 336, taken from EBI Alphafold repository (27). The catalytic residue (C264) shown in stick form with SH sidechain coloured yellow. (**B**) Surface representation of Ldt_Go2_, demonstrating flat bowl-like active site surrounding C264 (yellow). (**C**) Superimposition of Ldt_Go2_ (grey) with *Vibrio cholerae* LdtA (RCSB entry 7AJO, unreleased, blue; bound reaction intermediate at C444 shown in stick form), reveals the relatively more closed/capped cleft of 3-3 cross-link forming enzymes and outlines that potential donor and acceptor substrates of Ldt_Go2_ will likely be less constrained.

The discovery of enzymes forming 1-3 cross-links reaffirmed that the catalytic reactions carried out by domains belonging to the YkuD family are very diverse. This work expands our knowledge on peptidoglycan polymerization and opens new avenues to study how remodelling contributes to the maintenance of cell envelope integrity (20). *Acetobacteraceae* (also called acetic acid bacteria) are important for the food industry and are key organisms involved in the production of vinegar (30). These organisms have a high capacity to oxidize ethanol as well as various sugars to form acetic acid and display resistance to high concentrations of acetic acid released into the fermentative medium. It is tempting to speculate that the formation of 1-3 cross-links contributes to the maintenance of cell envelope integrity in these harsh conditions.

### Experimental procedures

#### Bacterial strains, plasmids, oligonucleotides and growth conditions

Bacterial strains, plasmids and oligonucleotides are described in Table S6. *G. oxydans* B58 (ATCC NRLL-BR8) and isogenic derivatives were grown in yeast peptone mannitol (YPM; 5 g/l yeast extract, 3 g/l peptone, 25 g/l mannitol) broth or agar at 30°C under agitation (200 rpm). *G. oxydans* cultures were inoculated with an overnight preculture at an OD_600nm_=0.05 and grown for 36h to stationary phase. *G. oxydans* transposon mutants were grown in the presence of kanamycin (100µg/ml) and gentamicin (10µg/ml) for complementation experiments Ldt_Go2_ expression in G. oxydans was induced by adding 100 ng/ml anhydrotetracycline to the media at an OD_600nm_=0.5. For heterologous expression, *E. coli* was grown in an auto-induction medium (31) at 30°C under agitation (200 rpm) supplemented with 100µg/ml ampicillin.

#### Plasmid constructions

Plasmid pBBR-TetR-Go2227 used to complement the transposon insertion in the *ldt_Go2_* is a derivative of pBBR1MCS-5-T_gdhM_*-tetR-mNG* allowing the inducible expression of proteins in *G. oxydans* under the control of the tetracycline promoter (25). pBBR-TetR-Go2227 was built using Golden Gate assembly. Three PCR fragments corresponding to (i) the gentamicin cassette (1632 bp), (ii) the pBBR1 origin of replication + the TetR gene (4029 bp), and (iii) the *ldt_Go2_*full length sequence (1021 bp) were amplified using oligos SM_0729 + SM_0730, SM_0725 + SM_0726 and SM_0727 + SM_0728, respectively using pBBR1MCS-5-T_gdhM_*-tetR-mNG* or *G. oxydans* chromosomal DNA as templates. The PCR products were purified by gel extraction, mixed in an equimolar ratio and assembled using the NEBridge^®^ Golden Gate Assembly Kit (BsaI-HF^®^ v2) according to manufacturer’s instructions. Recombinant plasmids were screened by PCR and plasmid candidates were fully sequenced by Plasmidsaurus (Plsamidsaurus.com) to confirm the absence of mutations. pET-Go2227, a pET2818 derivative expressing the full length Ldt_Go2_ enzyme was built using a synthetic DNA fragment with optimized codon usage for *E. coli* provided by Genewiz. The synthetic open reading frame corresponding to the full length Ldt_Go2_ gene (with a stop codon) was cloned into pET2818 as a NcoI-XhoI fragment.

### Preparation of *Gluconobacter oxydans* competent cells and transformation

*G. oxydans* was grown in 100 ml of YPM to an OD_600nm_ of 0.9 and spun for 10 min at 4,000 x g at 4°C. After 3 washes in 1mM HEPES buffer (pH7.0), cells were resuspended in 250 µl. Electroporation was carried out in 1 mm cuvettes using 50 µl of electrocompetent cells and 100 ng of plasmid in a volume of 1-2µl; parameters for electroporation were 2 kV, 25 µF and 200 Ω. After the pulse, 800µl of YPM media supplemented with 0.25% (m/v) MgSO_4_ and 0.15% (m/v) CaCl_2_ was added to the cells that were left to recover under agitation for 16 h before plating on YPM media supplemented with kanamycin and gentamicin.

#### Peptidoglycan extraction

*G. oxydans* and *E. coli* strains were grown until stationary phase in YPM or auto-induction medium, respectively. Cells were spun, supernatant discarded, and cell pellet snap frozen in liquid nitrogen. The cell pellet was resuspended in 20 ml of boiling MilliQ water (MQ) before the addition of SDS at a final concentration of 4% (m/v). After 30 min at 100 °C, the cells were cooled down to room temperature. Peptidoglycan was pelleted at 150,000 *g* for 1 h, washed five times using warm MQ water, freeze-dried and resuspended at a final concentration of 10 mg/ml.

#### Preparation of soluble muropeptides

2 mg of purified peptidoglycan was digested for 16h in 20 mM phosphate buffer (pH 5.5) supplemented with 250 Units of mutanolysin (Sigma) in a final volume of 200 μl. Following heat inactivation of mutanolysin (5 min at 100 °C), soluble disaccharide peptides were mixed with an equal volume of 250 mM borate buffer (pH 9.25) and reduced with 0.2 % (m/v) sodium borohydride. After 20 min at room temperature, the pH was adjusted to 5.0 using phosphoric acid. Reduced muropeptides were analyzed by RP-HPLC using a C18 analytical column (Hypersil Gold aQ, 1.9 µm particles, 150 × 2.1 mm; Thermo Fisher Scientific) at a temperature of 50°C. Muropeptide elution was performed at 0.3ml/min by applying a mixture of solvent A (water, 0.1% [v/v] formic acid) and solvent B (acetonitrile, 0.1% [v/v] formic acid). LC conditions were 0–12.5% B for 25 min increasing to 20% B for 10 min. After 5 min at 95%, the column was re-equilibrated for 10 min with 100% buffer A. UV absorbance at 202 nm was used to check the quality of samples and determine the volume to inject for LC-MS. A volume of sample with an intensity of the most abundant monomer of 1500mAU was used, giving an ion intensity of approximately 5.10^9^.

#### UHPLC-tandem mass spectrometry

An Ultimate 3000 Ultra High-Performance Chromatography (UHPLC; Dionex/Thermo Fisher Scientific) system coupled with a high-resolution Q Exactive Focus mass spectrometer (Thermo Fisher Scientific) was used for LC-HRMS analysis. Muropeptides were separated using a C18 analytical column (Hypersil Gold aQ, 1.9 µm particles, 150 × 2.1 mm; Thermo Fisher Scientific) at a temperature of 50°C. Muropeptide elution was performed as described in the previous paragraph. The Orbitrap Exploris 240 was operated under electrospray ionization (H-ESI high flow)-positive mode, full scan (m/z 150–2250) at resolution 120,000 (FWHM) at *m/z* 200, with normalized AGC Target 100%, and automated maximum ion injection time (IT). Data-dependent MS/MS were acquired on a ‘Top 5’ data-dependent mode using the following parameters: resolution 30,000; AGC 100%, automated IT, with normalized collision energy 25%.

#### Comparative genomics and bioinformatic analysis

Reference genomes and protein sequences (Table S5) were downloaded from NCBI Datasets (v15.25.0), and protein sequences were annotated locally using InterProScan (v5.64-96.0) (32,33). A custom Julia (34) script was then used to scrape the produced GFF3 files for YkuD-containing proteins and to extract their catalytic domains. Ldt_Go1_ and Ldt_Go2_ homologues were located by running a PSI-BLAST on the RefSeq Select database restricted to taxa in (Table S5) and iterating until no new hits were returned (26). Extracted YkuD proteins and Ldt_Go1/2_ homologues were aligned using Muscle (v5.1) (35), and maximum likelihood trees were constructed using IQ-TREE (v2.2.2.7) with ModelFinder (which selected WAG+R7 for Fig. 6A and WAG+F+G4 for Fig. 6B) and 1000 UFBoot replicates enabled (36–38). Trees were visualised and annotated using iTOL (v6.8.1) (39) with finishing touches applied in Inkscape (v1.3). ColabFold’s AlphaFold2_batch notebook (v1.5.2) was used with the default settings and relaxation enabled to obtain predicted structures for Ldt_Go1_, Ldt_Go2_, and their respective catalytic domains (27,40). Finally, Foldseek (v8-ef4e960) was used to search the AFDB50 database for structural homologues of Ldt_Go2_ (27,41).

## Data availability

LC-MS/MS datasets have been deposited in the GLYCOPOST repository (GPST000377). All plasmid sequences are available upon request. *G. oxydans* NRRL B58 genome has been deposited at DDBJ/ENA/GenBank under the accession <u>JAIPVW000000000</u>.

## Code availability

The script for PGFinder v1.1.0 is available at https://github.com/Mesnage-Org.

## Supporting information

This article contains supporting information (Fig. S1-S4, Table S1-S5 and Supplementary files S1-2).

## Acknowledgements

The authors thank Tino Pollen (Jülich Research Institute) for the gift of pBBR1-TetR plasmid to carry out complementation experiments. SM and ALL were supported by an MRC grant (MR/S009272/1). MGAZ is funded by a Mexican government PhD scholarship (CONAHCYT, 2021-000007-01EXTF-00221). BJR is the recipient of a NERC PhD studentship (NERC ACCE NE/S00713X/1). The work in BB’s lab was supported by Cornell University startup funds, an Academic Venture Fund award from the Atkinson Center for Sustainability at Cornell University, a Career Award at the Scientific Interface from the Burroughs Welcome Fund to BMB, ARPA-E award DE-AR0001341 to BB, and a gift from Mary Fernando Conrad and Tony Conrad to BMB.

## References

1. Weidel, W., and Pelzer, H. (1964) Bagshaped macromolecules - a new outlook on bacterial cell walls. Adv Enzymol Relat Areas Mol Biol 26, 193–232

2. Vollmer, W., Blanot, D., and de Pedro, M. A. (2008) Peptidoglycan structure and architecture. FEMS Microbiol Rev 32, 149–167

3. Ciak, J., and Hahn, F. E. (1957) Penicillin-induced lysis of *Escherichia coli*. Science 125, 119–120

4. Park, J. T., and Strominger, J. L. (1957) Mode of action of penicillin. Science 125, 99–101

5. Tipper, D. J., and Strominger, J. L. (1965) Mechanism of action of penicillins: a proposal based on their structural similarity to acyl-D-alanyl-D-alanine. Proc Natl Acad Sci U S A 54, 1133–1141

6. Cho, H., Uehara, T., and Bernhardt, T. G. (2014) Beta-lactam antibiotics induce a lethal malfunctioning of the bacterial cell wall synthesis machinery. Cell 159, 1300–1311

7. Wietzerbin, J., Das, B. C., Petit, J. F., Lederer, E., Leyh-Bouille, M., and Ghuysen, J. M. (1974) Occurrence of D-alanyl-(D)-meso-diaminopimelic acid and meso-diaminopimelyl-meso-diaminopimelic acid interpeptide linkages in the peptidoglycan of Mycobacteria. Biochemistry 13, 3471–3476

8. Lavollay, M., Arthur, M., Fourgeaud, M., Dubost, L., Marie, A., Veziris, N., Blanot, D., Gutmann, L., and Mainardi, J. L. (2008) The peptidoglycan of stationary-phase *Mycobacterium tuberculosis* predominantly contains cross-links generated by L,D-transpeptidation. J Bacteriol 190, 4360–4366

9. Mahapatra, S., Crick, D. C., McNeil, M. R., and Brennan, P. J. (2008) Unique structural features of the peptidoglycan of *Mycobacterium leprae*. J Bacteriol 190, 655–661

10. Peltier, J., Courtin, P., El Meouche, I., Lemee, L., Chapot-Chartier, M. P., and Pons, J. L. (2011) *Clostridium difficile* has an original peptidoglycan structure with a high level of N-acetylglucosamine deacetylation and mainly 3-3 cross-links. The Journal of biological chemistry 286, 29053–29062

11. Mainardi, J. L., Fourgeaud, M., Hugonnet, J. E., Dubost, L., Brouard, J. P., Ouazzani, J., Rice, L. B., Gutmann, L., and Arthur, M. (2005) A novel peptidoglycan cross-linking enzyme for a beta-lactam-resistant transpeptidation pathway. The Journal of biological chemistry 280, 38146–38152

12. Kim, H. S., Im, H. N., An, D. R., Yoon, J. Y., Jang, J. Y., Mobashery, S., Hesek, D., Lee, M., Yoo, J., Cui, M., Choi, S., Kim, C., Lee, N. K., Kim, S. J., Kim, J. Y., Bang, G., Han, B. W., Lee, B. I., Yoon, H. J., and Suh, S. W. (2015) The Cell Shape-determining Csd6 Protein from *Helicobacter pylori* Constitutes a New Family of L,D-Carboxypeptidase. The Journal of biological chemistry 290, 25103–25117

13. Godessart, P., Lannoy, A., Dieu, M., Van der Verren, S. E., Soumillion, P., Collet, J. F., Remaut, H., Renard, P., and De Bolle, X. (2021) beta-Barrels covalently link peptidoglycan and the outer membrane in the alpha-proteobacterium *Brucella abortus*. Nat Microbiol 6, 27–33

14. Sandoz, K. M., Moore, R. A., Beare, P. A., Patel, A. V., Smith, R. E., Bern, M., Hwang, H., Cooper, C. J., Priola, S. A., Parks, J. M., Gumbart, J. C., Mesnage, S., and Heinzen, R. A. (2021) beta-Barrel proteins tether the outer membrane in many Gram-negative bacteria. Nat Microbiol 6, 19–26

15. Bahadur, R., Chodisetti, P. K., and Reddy, M. (2021) Cleavage of Braun’s lipoprotein Lpp from the bacterial peptidoglycan by a paralog of l,d-transpeptidases, LdtF. Proc Natl Acad Sci U S A 118

16. Galley, N. F., Greetham, D., Alamán-Zárate, M.G., Williamson, M.P., Evans, C.A., Spittal, W.D., Buddle, J.E., Freeman, J., Davis, G.D., Dickman, M.J., Wilcox, M.H., Lovering, A.L., Fagan, R.P., Mesnage, S. Clostridioides difficile canonical L,D-transpeptidases catalyse a novel type of peptidoglycan cross-links and are not required for beta-lactam resistance Submitted for publication

17. Winkle, M., Hernandez-Rocamora, V. M., Pullela, K., Goodall, E. C. A., Martorana, A. M., Gray, J., Henderson, I. R., Polissi, A., and Vollmer, W. (2021) DpaA Detaches Braun’s Lipoprotein from Peptidoglycan. mBio 12, e00836–00821

18. Bern, M., Beniston, R., and Mesnage, S. (2017) Towards an automated analysis of bacterial peptidoglycan structure. Anal Bioanal Chem 409, 551–560

19. Sutterlin, L., Edoo, Z., Hugonnet, J. E., Mainardi, J. L., and Arthur, M. (2018) Peptidoglycan Cross-Linking Activity of L,D-Transpeptidases from *Clostridium difficile* and Inactivation of These Enzymes by beta-Lactams. Antimicrob Agents Chemother 62, e01607–01617

20. More, N., Martorana, A. M., Biboy, J., Otten, C., Winkle, M., Serrano, C. K. G., Monton Silva, A., Atkinson, L., Yau, H., Breukink, E., den Blaauwen, T., Vollmer, W., and Polissi, A. (2019) Peptidoglycan remodeling enables *Escherichia coli* to survive severe outer membrane assembly defect. mBio 10

21. Hernandez, S. B., Castanheira, S., Pucciarelli, M. G., Cestero, J. J., Rico-Perez, G., Paradela, A., Ayala, J. A., Velazquez, S., San-Felix, A., Cava, F., and Garcia-Del Portillo, F. (2022) Peptidoglycan editing in non-proliferating intracellular *Salmonella* as source of interference with immune signaling. PLoS Pathog 18, e1010241

22. Espaillat, A., Forsmo, O., El Biari, K., Bjork, R., Lemaitre, B., Trygg, J., Canada, F. J., de Pedro, M. A., and Cava, F. (2016) Chemometric analysis of bacterial peptidoglycan reveals atypical modifications that empower the cell wall against predatory enzymes and fly innate immunity. J Am Chem Soc 138, 9193–9204

23. Patel, A. V., Turner, R. D., Rifflet, A., Acosta-Martin, A. E., Nichols, A., Awad, M. M., Lyras, D., Gomperts-Boneca, I., Bern, M., Collins, M. O., and Mesnage, S. (2021) PGFinder, a novel analysis pipeline for the consistent, reproducible and high-resolution structural analysis of bacterial peptidoglycans. eLife 0:e70597

24. Schmitz, A. M., Pian, B., Medin, S., Reid, M. C., Wu, M., Gazel, E., and Barstow, B. (2021) Generation of a *Gluconobacter oxydans* knockout collection for improved extraction of rare earth elements. Nat Commun 12, 6693

25. Fricke, P. M., Lurkens, M., Hunnefeld, M., Sonntag, C. K., Bott, M., Davari, M. D., and Polen, T. (2021) Highly tunable TetR-dependent target gene expression in the acetic acid bacterium *Gluconobacter oxydans*. Appl Microbiol Biotechnol 105, 6835–6852

26. Camacho, C., Coulouris, G., Avagyan, V., Ma, N., Papadopoulos, J., Bealer, K., and Madden, T. L. (2009) BLAST+: architecture and applications. BMC Bioinformatics 10, 421

27. Jumper, J., Evans, R., Pritzel, A., Green, T., Figurnov, M., Ronneberger, O., Tunyasuvunakool, K., Bates, R., Zidek, A., Potapenko, A., Bridgland, A., Meyer, C., Kohl, S. A. A., Ballard, A. J., Cowie, A., Romera-Paredes, B., Nikolov, S., Jain, R., Adler, J., Back, T., Petersen, S., Reiman, D., Clancy, E., Zielinski, M., Steinegger, M., Pacholska, M., Berghammer, T., Bodenstein, S., Silver, D., Vinyals, O., Senior, A. W., Kavukcuoglu, K., Kohli, P., and Hassabis, D. (2021) Highly accurate protein structure prediction with AlphaFold. Nature 596, 583–589

28. van Kempen, M., Kim, S. S., Tumescheit, C., Mirdita, M., Lee, J., Gilchrist, C. L. M., Soding, J., and Steinegger, M. (2023) Fast and accurate protein structure search with Foldseek. Nat Biotechnol

29. Sacco, E., Hugonnet, J. E., Josseaume, N., Cremniter, J., Dubost, L., Marie, A., Patin, D., Blanot, D., Rice, L. B., Mainardi, J. L., and Arthur, M. (2010) Activation of the L,D-transpeptidation peptidoglycan cross-linking pathway by a metallo-D,D-carboxypeptidase in *Enterococcus faecium*. Mol Microbiol 75, 874–885

30. Gomes, R. J., Borges, M. F., Rosa, M. F., Castro-Gomez, R. J. H., and Spinosa, W. A. (2018) Acetic Acid Bacteria in the Food Industry: Systematics, Characteristics and Applications. Food Technol Biotechnol 56, 139–151

31. Studier, F. W. (2018) T7 expression systems for inducible production of proteins from cloned genes in *Escherichia coli*. Curr Protoc Mol Biol 124, e63

32. Blum, M., Chang, H. Y., Chuguransky, S., Grego, T., Kandasaamy, S., Mitchell, A., Nuka, G., Paysan-Lafosse, T., Qureshi, M., Raj, S., Richardson, L., Salazar, G. A., Williams, L., Bork, P., Bridge, A., Gough, J., Haft, D. H., Letunic, I., Marchler-Bauer, A., Mi, H., Natale, D. A., Necci, M., Orengo, C. A., Pandurangan, A. P., Rivoire, C., Sigrist, C. J. A., Sillitoe, I., Thanki, N., Thomas, P. D., Tosatto, S. C. E., Wu, C. H., Bateman, A., and Finn, R. D. (2021) The InterPro protein families and domains database: 20 years on. Nucleic Acids Res 49, D344–D354

33. Jones, P., Binns, D., Chang, H. Y., Fraser, M., Li, W., McAnulla, C., McWilliam, H., Maslen, J., Mitchell, A., Nuka, G., Pesseat, S., Quinn, A. F., Sangrador-Vegas, A., Scheremetjew, M., Yong, S. Y., Lopez, R., and Hunter, S. (2014) InterProScan 5: genome-scale protein function classification. Bioinformatics 30, 1236–1240

34. Bezanson, J., Edelman, A., Karpinski, S., and Shah, V. B. (2017) Julia: a fresh approach to numerical computing. SIAM Reviews 59, 65–98

35. Edgar, R. C. (2004) MUSCLE: multiple sequence alignment with high accuracy and high throughput. Nucleic Acids Res 32, 1792–1797

36. Hoang, D. T., Chernomor, O., von Haeseler, A., Minh, B. Q., and Vinh, L. S. (2018) UFBoot2: Improving the Ultrafast Bootstrap Approximation. Mol Biol Evol 35, 518–522

37. Kalyaanamoorthy, S., Minh, B. Q., Wong, T. K. F., von Haeseler, A., and Jermiin, L. S. (2017) ModelFinder: fast model selection for accurate phylogenetic estimates. Nat Methods 14, 587–589

38. Minh, B. Q., Schmidt, H. A., Chernomor, O., Schrempf, D., Woodhams, M. D., von Haeseler, A., and Lanfear, R. (2020) IQ-TREE 2: New Models and Efficient Methods for Phylogenetic Inference in the Genomic Era. Mol Biol Evol 37, 1530–1534

39. Letunic, I., and Bork, P. (2021) Interactive Tree Of Life (iTOL) v5: an online tool for phylogenetic tree display and annotation. Nucleic Acids Res 49, W293–W296

40. Mirdita, M., Schutze, K., Moriwaki, Y., Heo, L., Ovchinnikov, S., and Steinegger, M. (2022) ColabFold: making protein folding accessible to all. Nat Methods 19, 679–682

41. Varadi, M., Anyango, S., Deshpande, M., Nair, S., Natassia, C., Yordanova, G., Yuan, D., Stroe, O., Wood, G., Laydon, A., Zidek, A., Green, T., Tunyasuvunakool, K., Petersen, S., Jumper, J., Clancy, E., Green, R., Vora, A., Lutfi, M., Figurnov, M., Cowie, A., Hobbs, N., Kohli, P., Kleywegt, G., Birney, E., Hassabis, D., and Velankar, S. (2022) AlphaFold Protein Structure Database: massively expanding the structural coverage of protein-sequence space with high-accuracy models. Nucleic Acids Res 50, D439–D444

42. Mistry, J., Chuguransky, S., Williams, L., Qureshi, M., Salazar, G. A., Sonnhammer, E. L. L., Tosatto, S. C. E., Paladin, L., Raj, S., Richardson, L. J., Finn, R. D., and Bateman, A. (2021) Pfam: The protein families database in 2021. Nucleic Acids Res 49, D412–D419

43. Wang, J., Chitsaz, F., Derbyshire, M. K., Gonzales, N. R., Gwadz, M., Lu, S., Marchler, G. H., Song, J. S., Thanki, N., Yamashita, R. A., Yang, M., Zhang, D., Zheng, C., Lanczycki, C. J., and Marchler-Bauer, A. (2023) The conserved domain database in 2023. Nucleic Acids Res 51, D384–D388

