## Supplementary Figures S1-S4 for "Unusual 1-3 peptidoglycan cross-links in *Acetobacteraceae* are made by L,D-transpeptidases with a catalytic domain distantly related to YkuD domains"

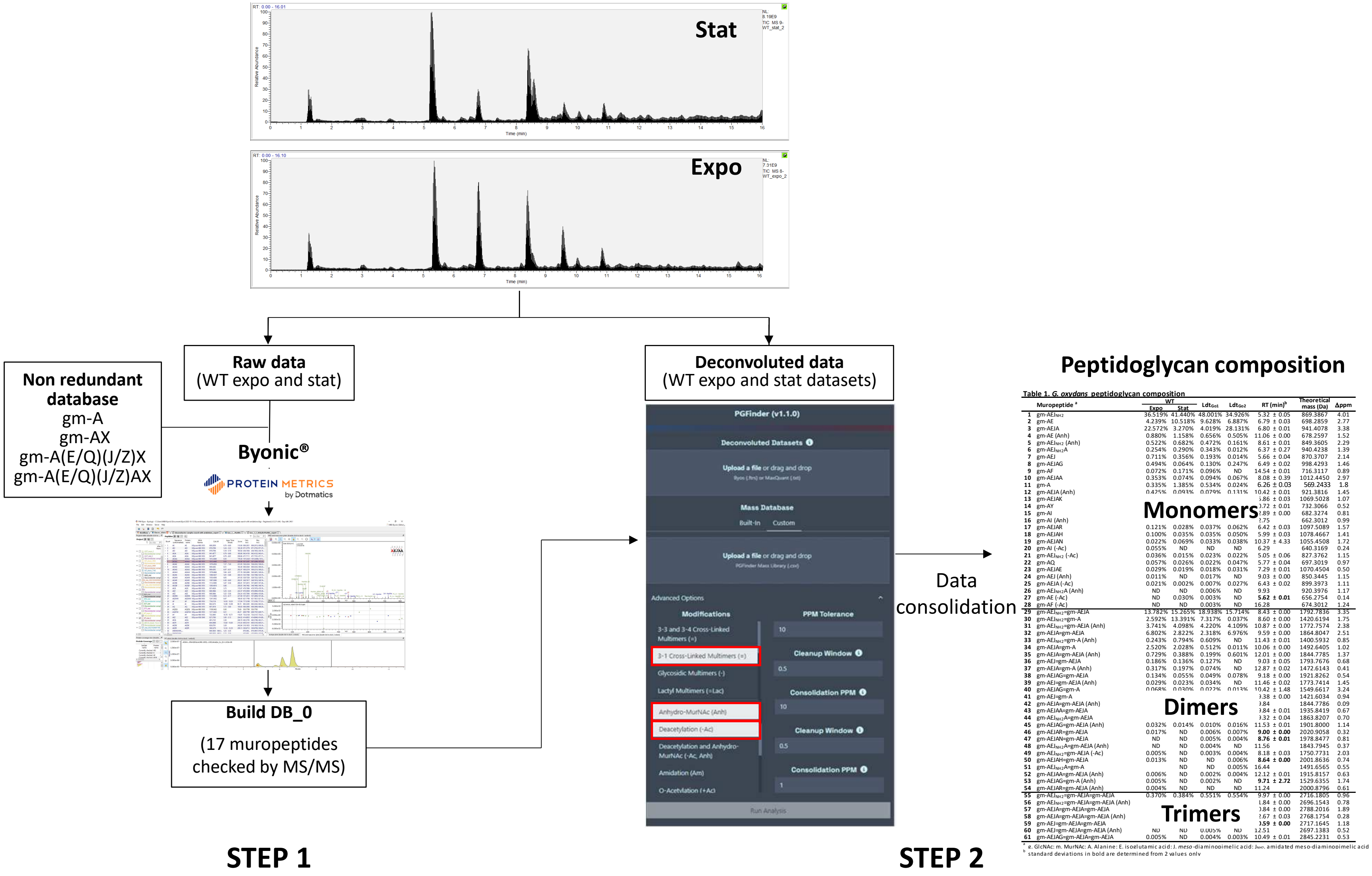


**Fig. S1. Strategy for *G. oxydans* peptidoglycan structural analysis.** A first search was performed using the Byonic™ module from the Byos® software to identify monomers based on MS/MS data. The search space contained disaccharide substituted by mono-, di-, tri-, tetra- and pentapeptides stems containing glutamic acid (E) or glutamine (Q) in position 2, *meso*-diaminopimelic acid (J) or amidated *meso*-diaminopimelic acid (Z) in position 3, any possible aminoacids (X) in position 4 or the AX dipeptides in positions 4 and 5. The 17 monomers identified by MS/MS were combined to generate the database DB_0. A second search was performed with PGFinder, enabling the formation of dynamic libraries containing dimers and trimers resulting from 4-3, 3-3 and 1-3 cross-links as well as their derivatives containing deacetylated or anhydroMurNAc sugar moeities. The corresponding options are boxes in red in the graphic user interface.

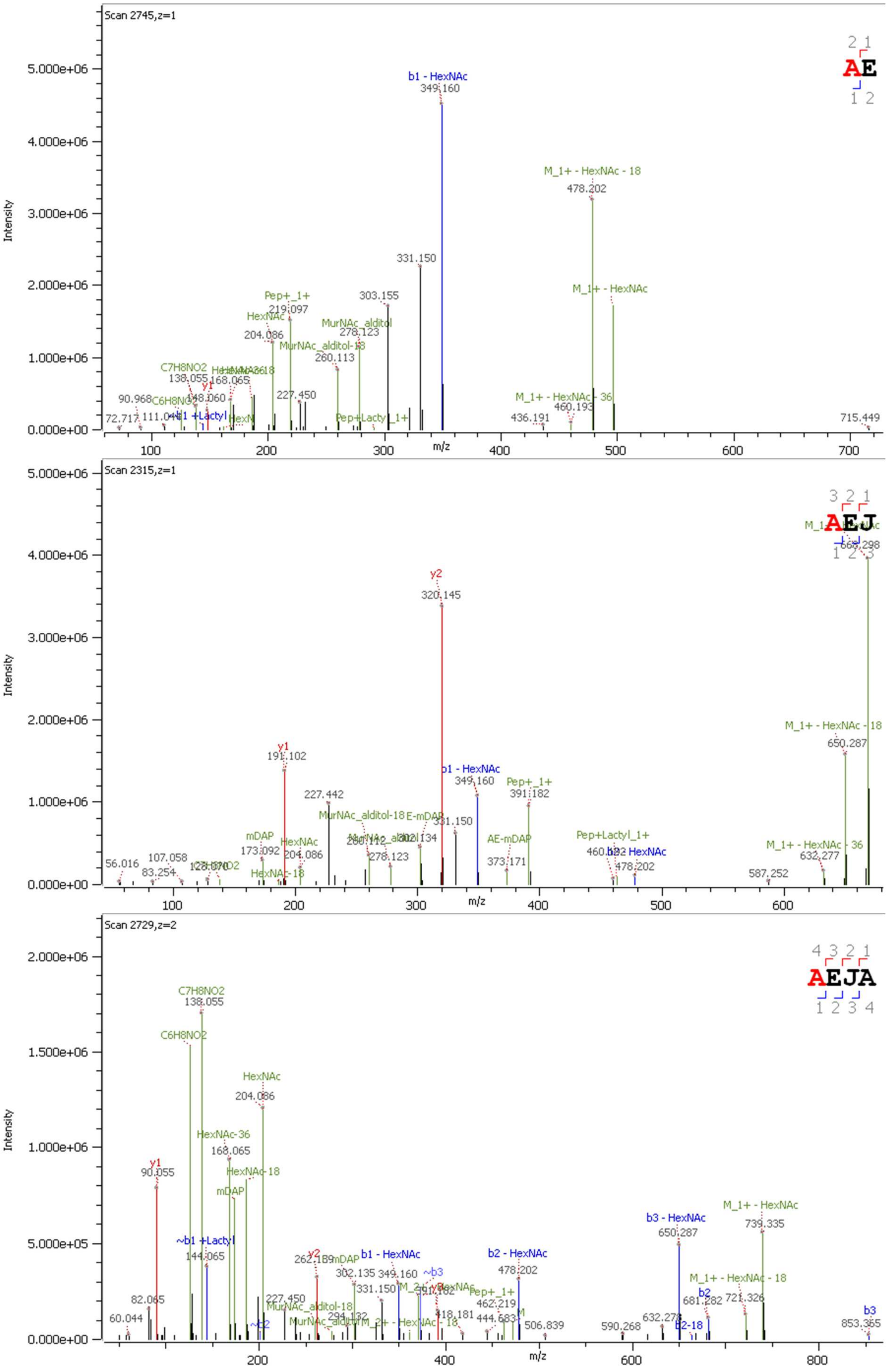

Scan 2745,z=1
Intensity
m/z
AE
b1 - HexNAc
349.160
M_1+ - HexNAc - 18
478.202
331.150
303.155
M_1+ - HexNAc
Pep+_1+
219.097
HexNAc
204.086
MurNAc_alditol
278.123
MurNAc_alditol-18
260.113
C7H8NO2
138.055
168.065
y1
148.060
90.968
C6H8NO2
227.450
72.717
111.046
b1 +Lactyl
Pep+Lactyl_1+
M_1+ - HexNAc - 36
460.193
436.191
715.449
Scan 2315,z=1
AEJ
667.298
y2
320.145
M_1+ - HexNAc - 18
650.287
y1
191.102
b1 - HexNAc
349.160
Pep+_1+
391.182
227.442
MurNAc_alditol-18
E-mDAP
331.150
302.134
mDAP
173.092
HexNAc
204.086
AE-mDAP
373.171
Pep+Lactyl_1+
b2 - HexNAc
478.202
M_1+ - HexNAc - 36
632.277
56.016
107.058
83.254
HexNAc-18
278.123
587.252
Scan 2729,z=2
AEJA
C7H8NO2
138.055
C6H8NO2
HexNAc
204.086
HexNAc-36
168.065
y1
90.055
HexNAc-18
mDAP
~b1 +Lactyl
144.065
y2
262.139
b1 - HexNAc
349.160
302.135
331.150
~b3
b2 - HexNAc
478.202
b3 - HexNAc
650.287
M_1+ - HexNAc
739.335
M_1+ - HexNAc - 18
721.326
b2
681.282
b3
853.365
82.065
60.044
227.450
MurNAc_alditol-18
294.132
Pep+_1+
462.219
418.181
444.683
506.839
590.268
632.278
b2-18

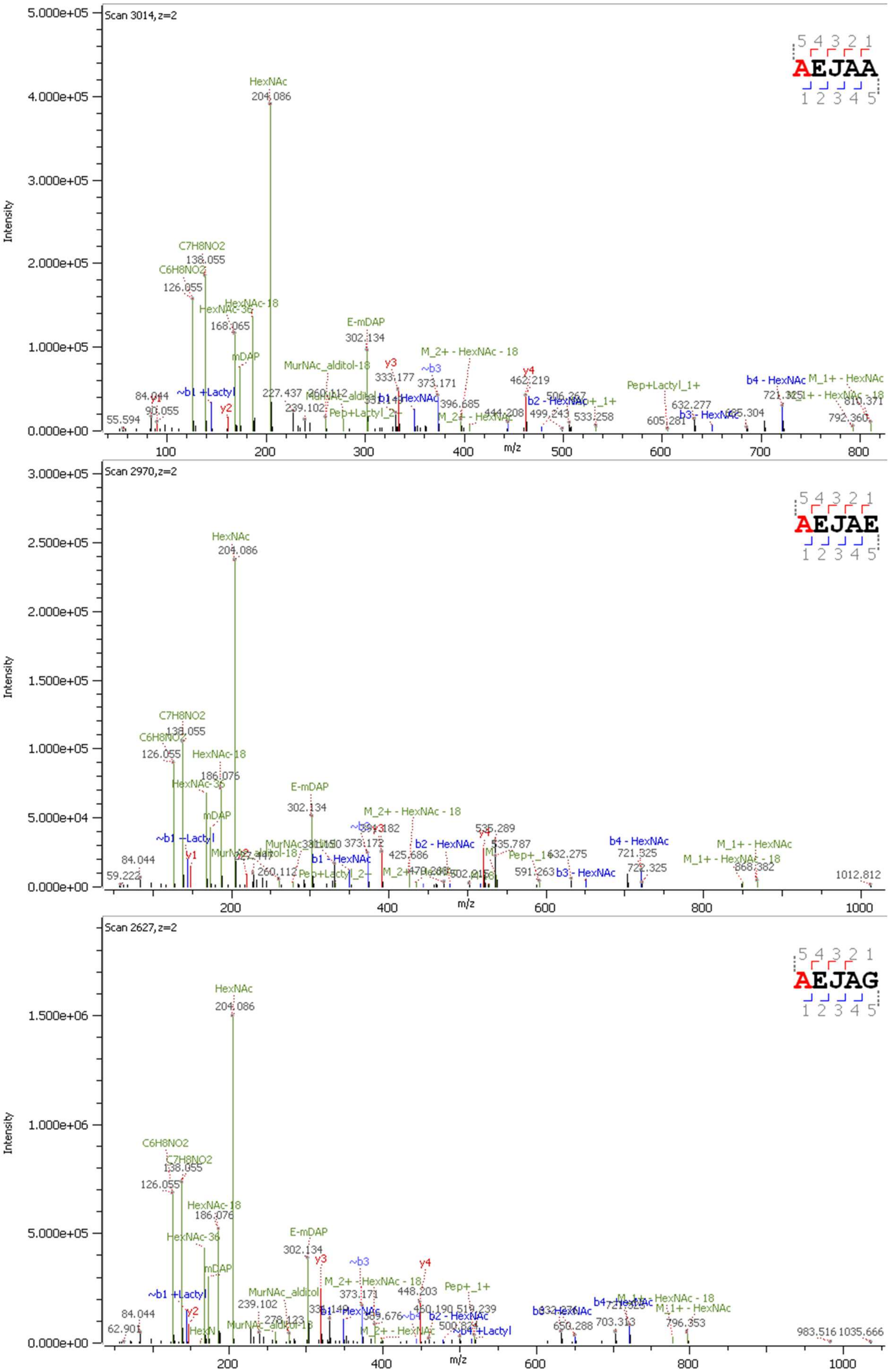

Scan 3014, z=2
AEJAA
5 4 3 2 1
1 2 3 4 5
Intensity
m/z
HexNAc
204.086
C7H8NO2
138.055
C6H8NO2
126.055
HexNAc-18
HexNAc-36
168.065
mDAP
E-mDAP
302.134
MurNAc_alditol-18
M_2+ - HexNAc - 18
Pep+Lactyl_1+
b4 - HexNAc
M_1+ - HexNAc
M_1+ - HexNAc - 18
~b1 +Lactyl
b2 - HexNAc
b3 - HexNAc
Scan 2970, z=2
AEJAE
186.076
535.289
721.525
1012.812
Scan 2627, z=2
AEJAG
MurNAc_alditol
Pep+_1+
703.313
796.353
983.516
1035.666

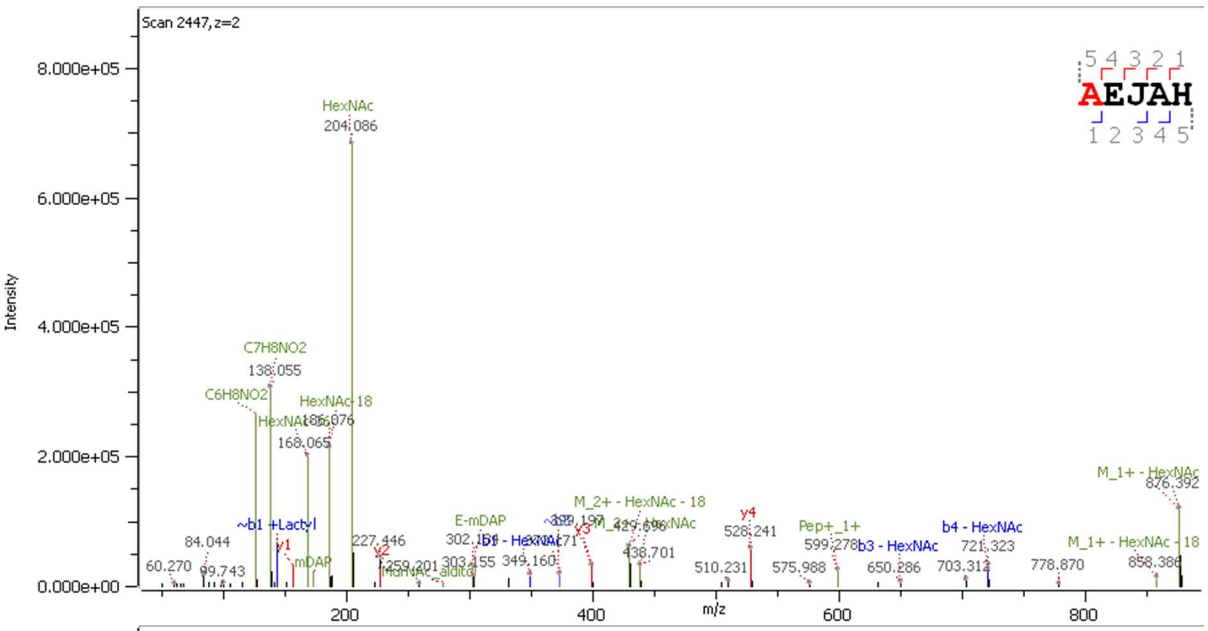
Scan 2447, z=2
Intensity
m/z
AEJAH
HexNAc
204.086
C7H8NO2
138.055
C6H8NO2
HexNAc-18
186.076
168.065
~b1 +Lactyl
y1
mDAP
y2
227.446
84.044
60.270
99.743
E-mDAP
302.154
303.155
349.160
y3
M_2+ - HexNAc - 18
438.701
y4
528.241
510.231
575.988
Pep+_1+
599.278
b3 - HexNAc
650.286
b4 - HexNAc
721.323
703.312
778.870
M_1+ - HexNAc
876.392
M_1+ - HexNAc - 18
858.386

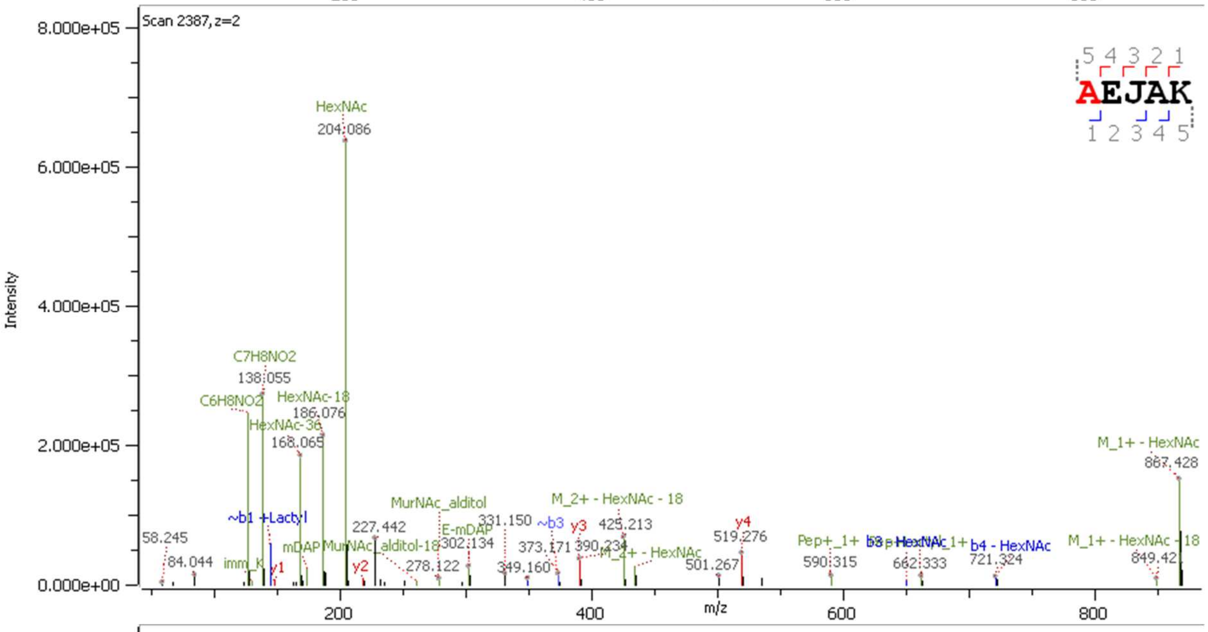
Scan 2387, z=2
Intensity
m/z
AEJAK
HexNAc
204.086
C7H8NO2
138.055
C6H8NO2
HexNAc-18
186.076
HexNAc-36
168.065
~b1 +Lactyl
58.245
84.044
imm_K
y1
mDAP
y2
227.442
MurNAc_alditol
MurNAc_alditol-18
278.122
E-mDAP
302.134
331.150
~b3
373.171
y3
390.234
425.213
M_2+ - HexNAc - 18
M_2+ - HexNAc
349.160
y4
519.276
501.267
Pep+_1+
590.315
662.333
b4 - HexNAc
721.324
M_1+ - HexNAc
867.428
M_1+ - HexNAc - 18
849.42

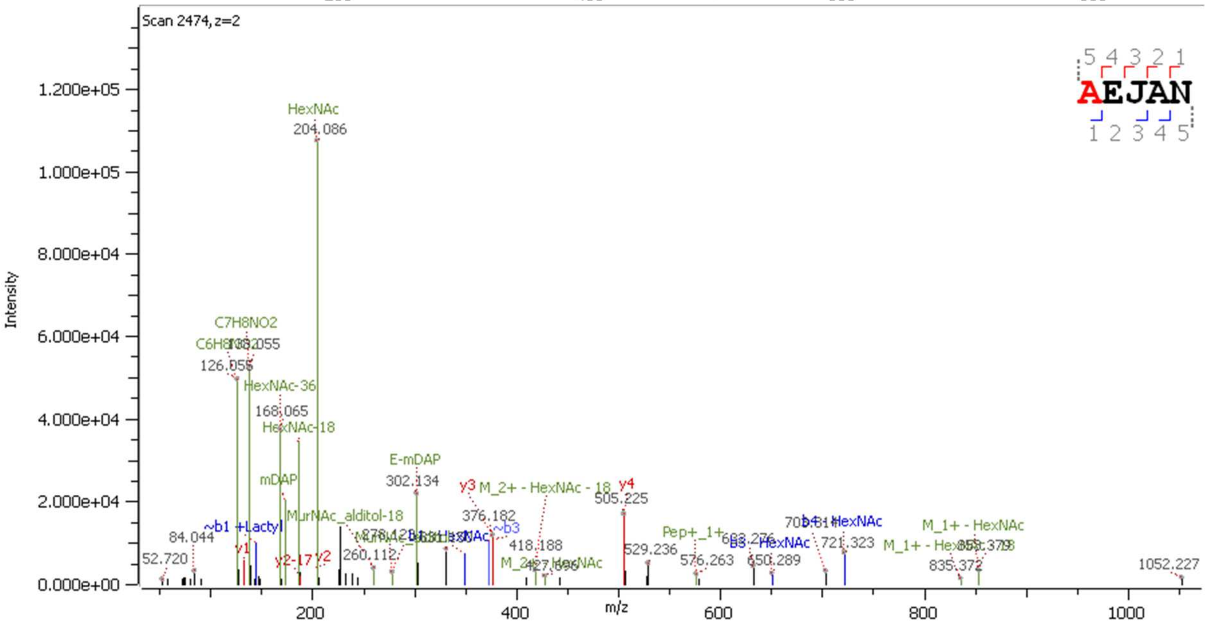
Scan 2474, z=2
Intensity
m/z
AEJAN
HexNAc
204.086
C7H8NO2
138.055
126.055
HexNAc-36
168.065
HexNAc-18
mDAP
~b1 +Lactyl
84.044
52.720
y1
y2-17
y2
MurNAc_alditol-18
260.112
E-mDAP
302.134
y3
M_2+ - HexNAc - 18
376.182
~b3
418.188
y4
505.225
529.236
Pep+_1+
576.263
b3 - HexNAc
650.289
b4 - HexNAc
721.323
M_1+ - HexNAc
835.372
1052.227

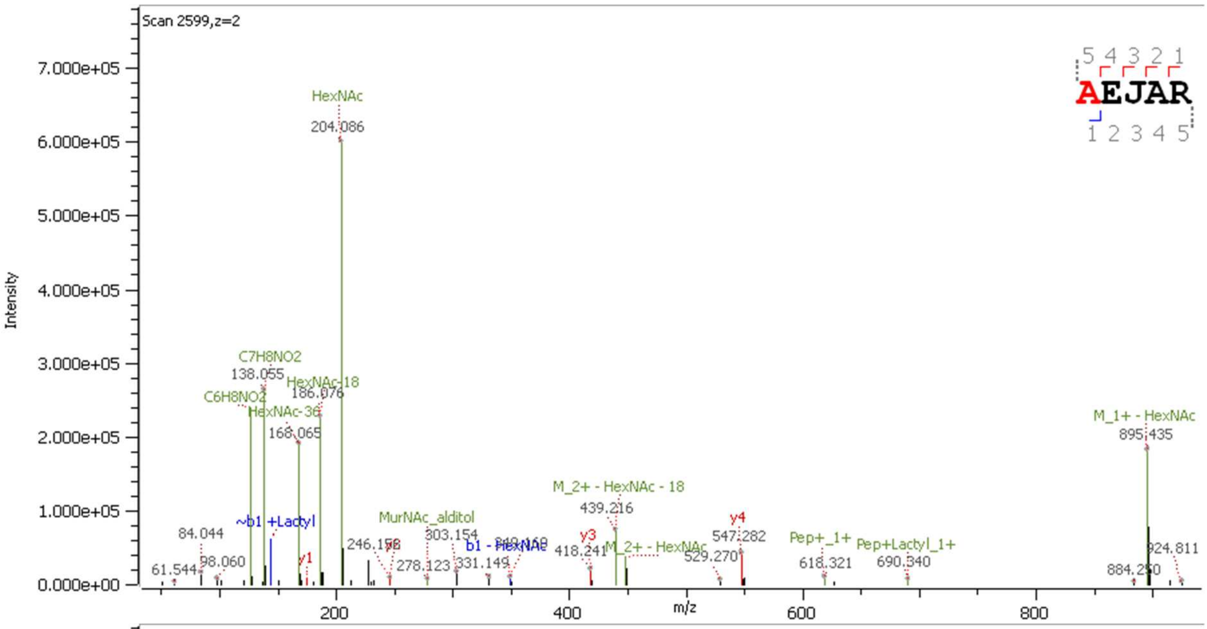
Scan 2599,z=2
Intensity
m/z
AEJAR
HexNAc
204.086
C7H8NO2
138.055
HexNAc-18
186.076
C6H8NO2
HexNAc-36
168.065
M_1+ - HexNAc
895.435
M_2+ - HexNAc - 18
439.216
~b1 +Lactyl
MurNAc_alditol
303.154
84.044
246.158
y4
547.282
Pep+_1+
Pep+Lactyl_1+
b1 - HexNAc
y3
418.241
M_2+ - HexNAc
924.811
61.544
98.060
y1
278.123
331.149
529.270
618.321
690.340
884.250

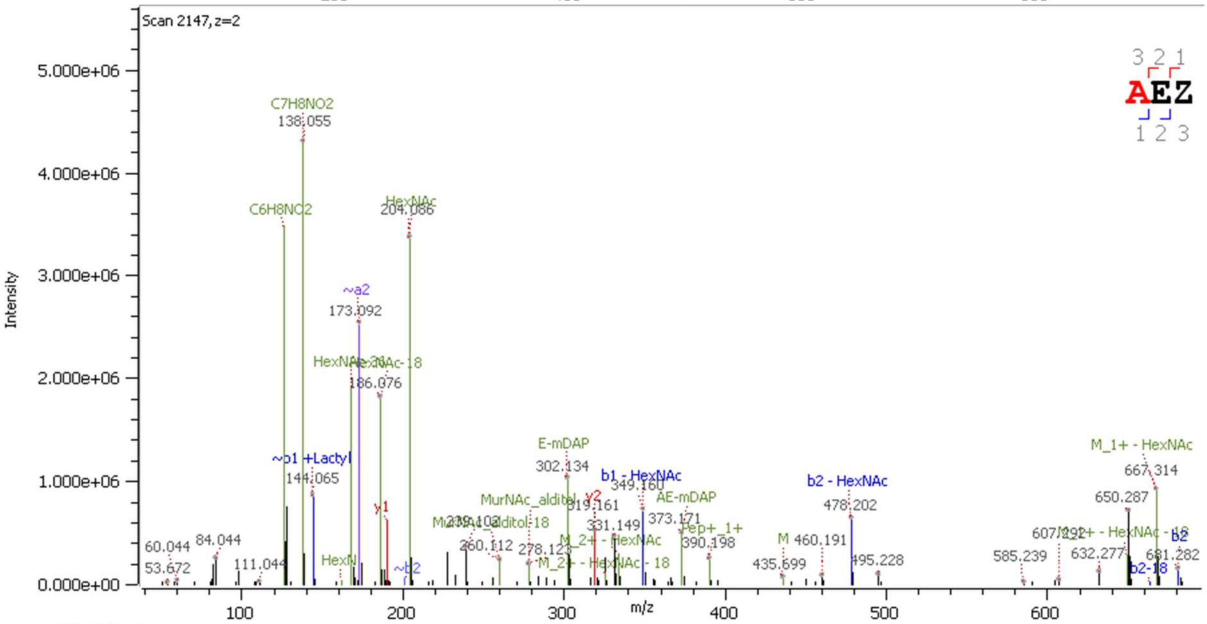
Scan 2147,z=2
Intensity
m/z
AEZ
C7H8NO2
138.055
C6H8NO2
HexNAc
204.086
~a2
173.092
HexNAc-18
186.076
M_1+ - HexNAc
667.314
E-mDAP
302.134
~b1 +Lactyl
144.065
b1 - HexNAc
349.160
b2 - HexNAc
478.202
AE-mDAP
373.171
650.287
MurNAc_alditol
y2
319.161
331.149
y1
Pep+_1+
390.198
460.191
60.044
84.044
278.123
260.112
M_2+ - HexNAc - 18
585.239
632.277
681.282
53.672
111.044
435.699
495.228
b2-18
b2

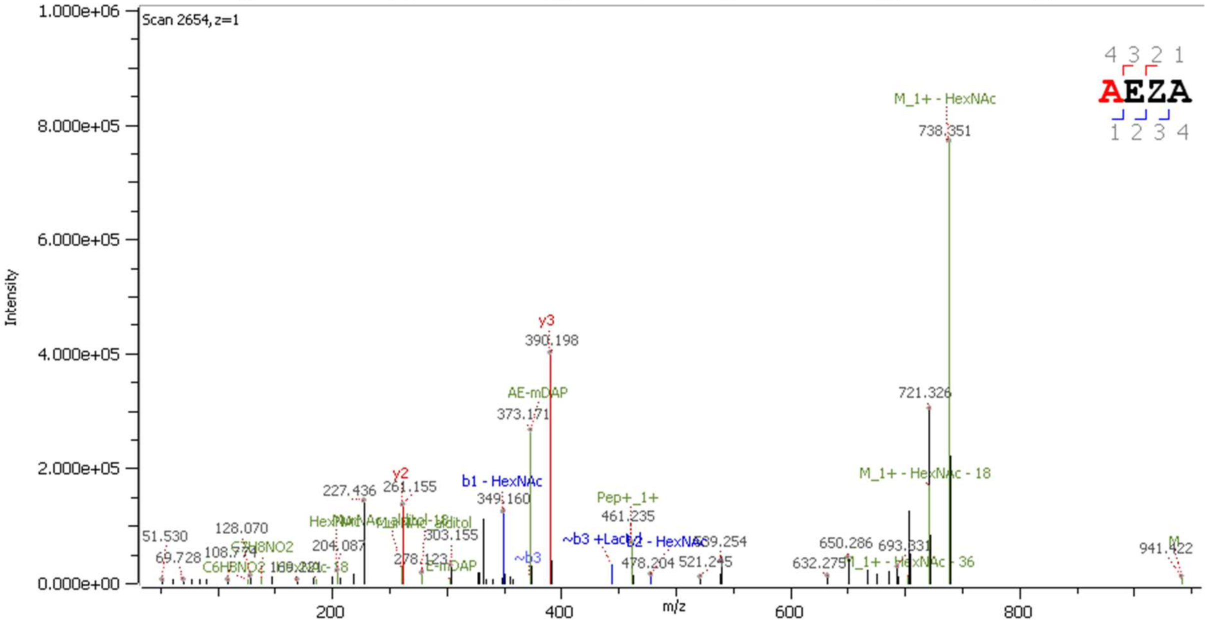
Scan 2654,z=1
Intensity
m/z
AEZA
M_1+ - HexNAc
738.351
y3
390.198
AE-mDAP
373.171
721.326
M_1+ - HexNAc - 18
y2
261.155
227.436
b1 - HexNAc
349.160
Pep+_1+
461.235
51.530
128.070
C7H8NO2
204.087
303.155
~b3 +Lactyl
b2 - HexNAc
539.254
650.286
693.331
941.422
69.728
108.774
C6H8NO2
278.123
E-mDAP
~b3
478.204
521.245
632.275
M_1+ - HexNAc - 36

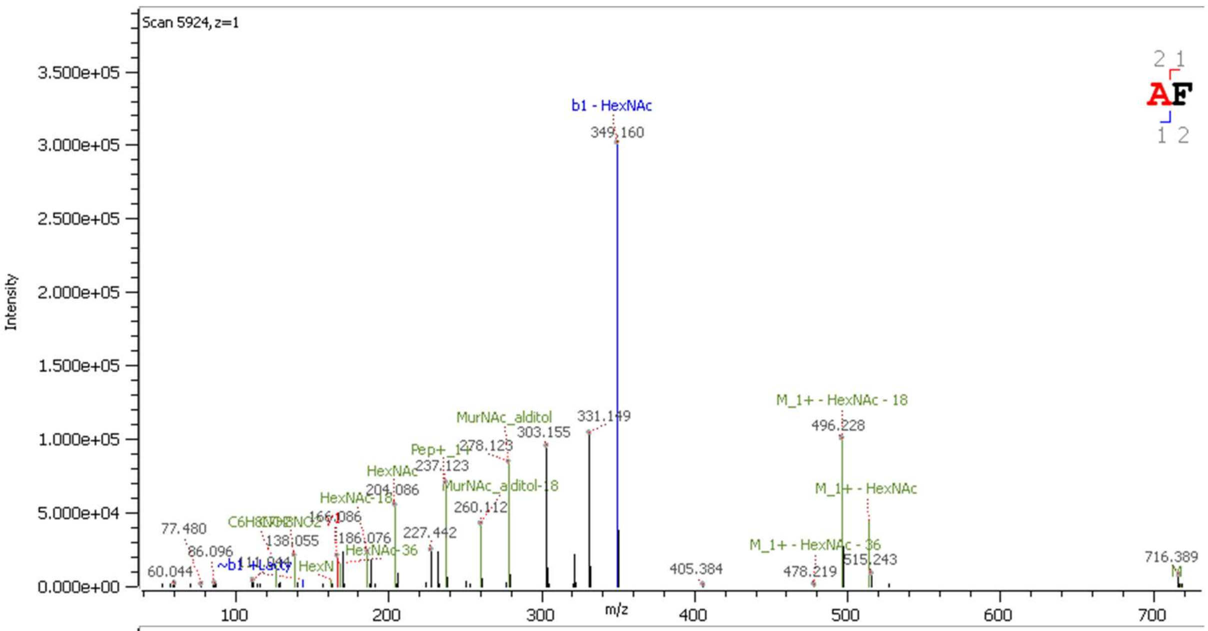
Scan 5924, z=1
Intensity
m/z
b1 - HexNAc
349.160
331.149
MurNAc_alditol
303.155
278.123
Pep+_1+
237.123
HexNAc
204.086
MurNAc_alditol-18
260.112
HexNAc-18
166.086
227.442
186.076
HexNAc-36
138.055
C6H8NO2
77.480
86.096
60.044
~b1 +Lactyl
M_1+ - HexNAc - 18
496.228
M_1+ - HexNAc
M_1+ - HexNAc - 36
515.243
478.219
405.384
716.389
M
AF

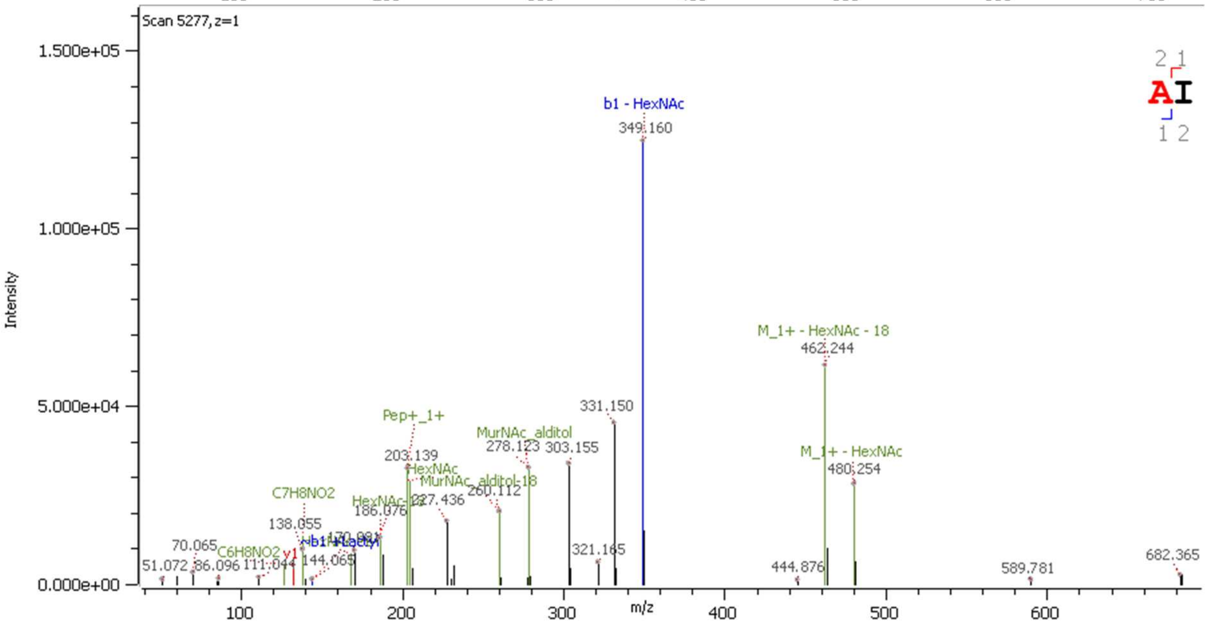
Scan 5277, z=1
Intensity
m/z
b1 - HexNAc
349.160
331.150
M_1+ - HexNAc - 18
462.244
M_1+ - HexNAc
480.254
Pep+_1+
203.139
HexNAc
MurNAc_alditol
278.123
303.155
MurNAc_alditol-18
227.436
260.112
HexNAc-18
186.076
C7H8NO2
138.055
C6H8NO2
y1
~b1 +Lactyl
170.091
144.065
70.065
51.072
86.096
111.044
321.165
444.876
589.781
682.365
AI

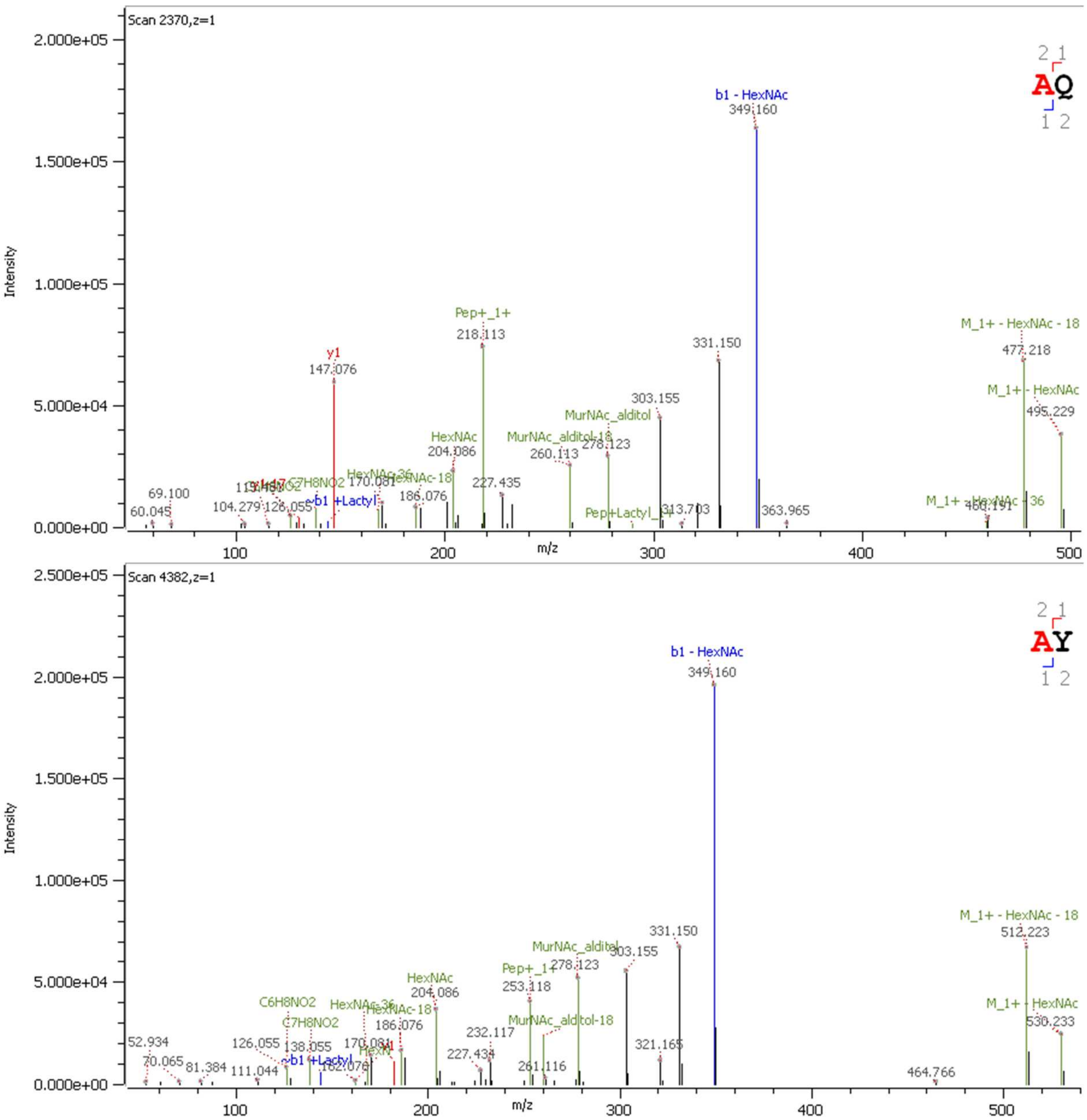


**Figure S2. Tandem mass spectrometry analysis of *G. oxydans* monomers using the Byonic™ module.**

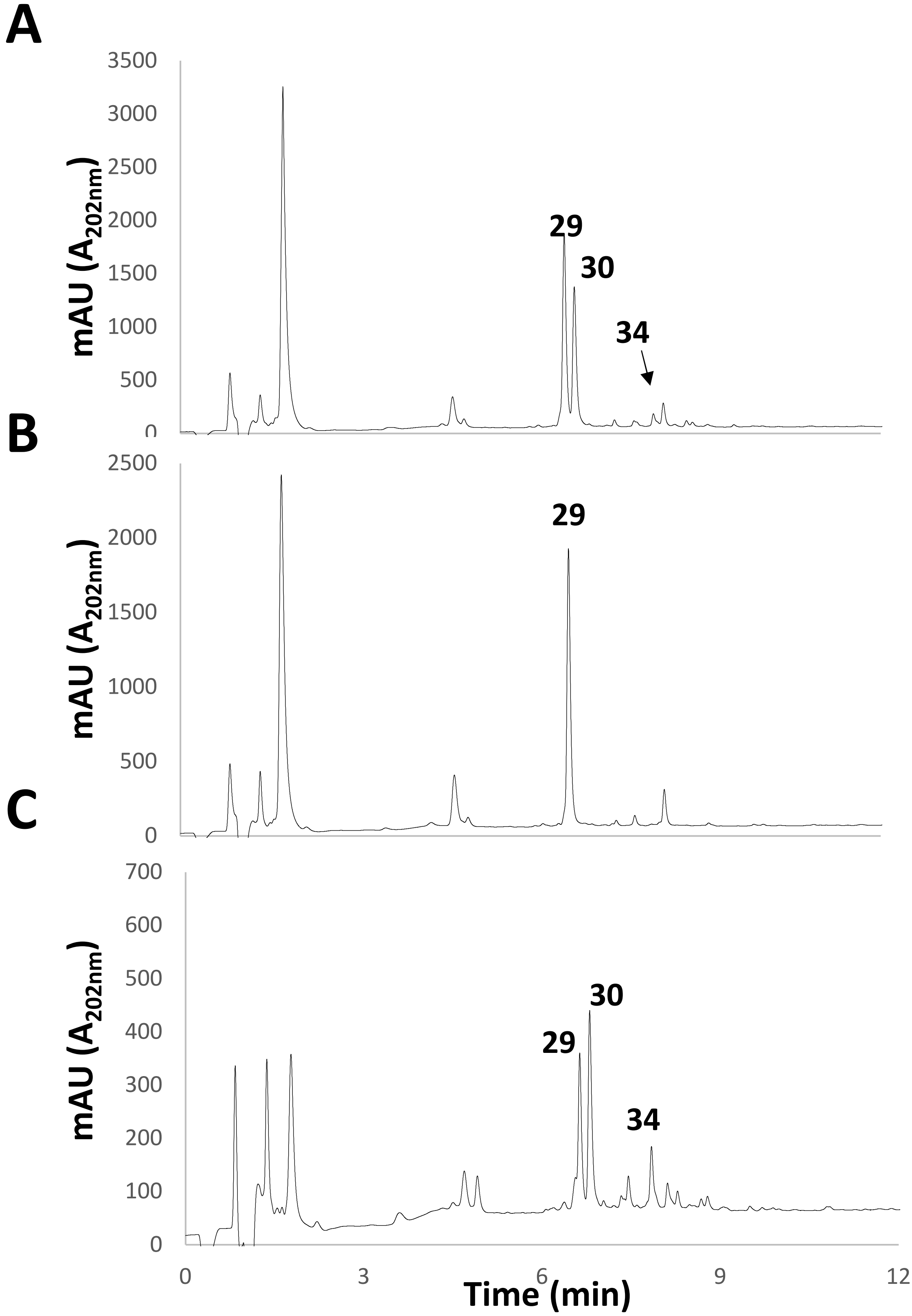


**Fig. S3. Complementation of the *ldt*$_{Go2}$ transposon insertion restores 1-3 cross-links.** Wild-type train B58 (**A**), the *ldtGo2* insertion mutant (**B**) and the complemented mutant (**C**) were grown in YPM media. The expression of *ldt*$_{Go2}$ in the complemented strain was induced at 0D600=0.5 with 100 ng/ml anhydrotetracycline. The peaks corresponding to the major species are labelled. Numbers refer to the muropeptides described in Table 1.

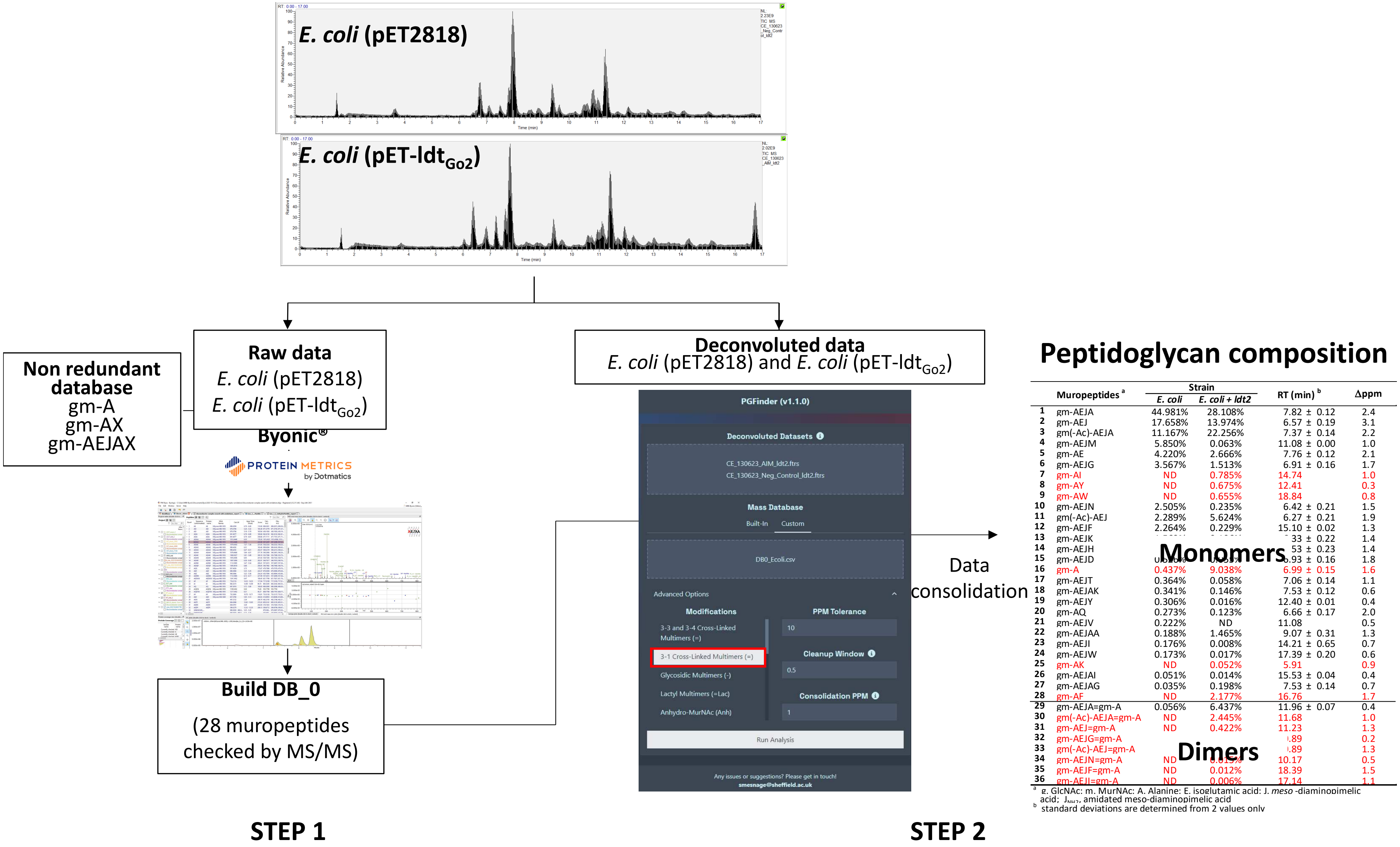


**Fig. S4. Strategy for peptidoglycan structural analysis of *E. coli* producing Ldt$_{Go2}$.** A first search was performed using the Byonic™ module from Byos® to identify monomers based on MS/MS data. The search space contained disaccharide substituted by mono-, di-, tri-, tetra- and pentapeptides stems containing glutamic acid (E) or glutamine (Q) in position 2, *meso*-diaminopimelic acid (J) or amidated *meso*-diaminopimelic acid (Z) in position 3, any possible aminoacids (X) in position 4 or the AX dipeptides in positions 4 and 5. The 28 monomers identified by MS/MS were combined to generate the database DB_0_Ec. A second search was performed with PGFinder, enabling the formation of dynamic libraries containing dimers and trimers resulting from 1-3 cross-links only (boxed in red).
