## Supplementary Tables S1-S6 for "Unusual 1-3 peptidoglycan cross-links in *Acetobacteraceae* are made by L,D-transpeptidases with a catalytic domain distantly related to YkuD domains"

### Table S1. List of peptide sequences used for *G. oxydans* MS/MS searches

| | |
|---|---|
| A | AEZAM |
| AE | AEZAN |
| AA | AEZAP |
| AC | AEZAQ |
| AR | AEZAR |
| AN | AEZAS |
| AD | AEZAT |
| AQ | AEZAV |
| AG | AEZAW |
| AH | AEZAY |
| AI | AQJ |
| AL | AQJA |
| AK | AQJC |
| AM | AQJD |
| AF | AQJE |
| AP | AQJF |
| AS | AQJG |
| AT | AQJH |
| AW | AQJI |
| AY | AQJK |
| AV | AQJM |
| AEJ | AQJN |
| AEJA | AQJP |
| AEJC | AQJQ |
| AEJD | AQJR |
| AEJE | AQJS |
| AEJF | AQJT |
| AEJG | AQJV |
| AEJH | AQJW |
| AEJI | AQJY |
| AEJK | AQJAA |
| AEJM | AQJAC |
| AEJN | AQJAD |
| AEJP | AQJAE |
| AEJQ | AQJAF |
| AEJR | AQJAG |
| AEJS | AQJAH |
| AEJT | AQJAI |
| AEJV | AQJAK |
| AEJW | AQJAM |
| AEJY | AQJAN |
| AEJAA | AQJAP |
| AEJAC | AQJAQ |
| AEJAD | AQJAR |
| AEJAE | AQJAS |
| AEJAF | AQJAT |
| AEJAG | AQJAV |
| AEJAH | AQJAW |
| AEJAI | AQJAY |
| AEJAK | AQZ |
| AEJAM | AQZA |
| AEJAN | AQZC |
| AEJAP | AQZD |
| AEJAQ | AQZE |
| AEJAR | AQZF |
| AEJAS | AQZG |
| AEJAT | AQZH |
| AEJAV | AQZI |
| AEJAW | AQZK |
| AEJAY | AQZM |
| AEZ | AQZN |
| AEZA | AQZP |
| AEZC | AQZQ |
| AEZD | AQZR |
| AEZE | AQZS |
| AEZF | AQZT |
| AEZG | AQZV |
| AEZH | AQZW |
| AEZI | AQZY |
| AEZK | AQZAA |
| AEZM | AQZAC |
| AEZN | AQZAD |
| AEZP | AQZAQ |
| AEZQ | AQZAF |
| AEZR | AQZAG |
| AEZS | AQZAH |
| AEZT | AQZAI |
| AEZV | AQZAK |
| AEZW | AQZAM |
| AEZY | AQZAN |
| AEZAA | AQZAP |
| AEZAC | AQZAQ |
| AEZAD | AQZAR |
| AEZAE | AQZAS |
| AEZAF | AQZAT |
| AEZAG | AQZAV |
| AEZAH | AQZAW |
| AEZAI | AQZAY |
| AEZAK | |

**Table S2. Database for *G. oxydans* PGFinder searches**

| Structure | Monoisotopicmass |
|---|---|
| gm-A\|1 | 569.24331 |
| gm-AE\|1 | 698.2859 |
| gm-AF\|1 | 716.31172 |
| gm-AI\|1 | 682.32737 |
| gm-AQ\|1 | 697.30189 |
| gm-AY\|1 | 732.30664 |
| gm-AEJ\|1 | 870.37069 |
| gm-AEZ\|1 | 869.38668 |
| gm-AEJA\|1 | 941.40783 |
| gm-AEZA\|1 | 940.42382 |
| gm-AEJAA\|1 | 1012.44497 |
| gm-AEJAE\|1 | 1070.45042 |
| gm-AEJAG\|1 | 998.42929 |
| gm-AEJAH\|1 | 1078.46674 |
| gm-AEJAK\|1 | 1069.50279 |
| gm-AEJAN\|1 | 1055.45076 |
| gm-AEJAR\|1 | 1097.50894 |

**Table S3. List of peptide sequences used for *E. coli* MS/MS searches**

A
AE
AC
AD
AF
AG
AH
AI
AJ
AK
AL
AM
AN
AP
AQ
AR
AS
AT
AV
AW
AY
AZ
AEJ
AEJA
AEJC
AEJD
AEJE
AEJF
AEJG
AEJH
AEJI
AEJK
AEJM
AEJN
AEJP
AEJQ
AEJR
AEJS
AEJT
AEJV
AEJW
AEJY
AEJAA
AEJAC
AEJAD
AEJAE
AEJAF
AEJAG
AEJAH
AEJAI
AEJAK
AEJAM
AEJAN
AEJAP
AEJAQ
AEJAR
AEJAS
AEJAT
AEJAV
AEJAW
AEJAY

**Table S2. Database for *E. coli* PGFinder searches**

| Structure | Monoisotopic mass |
|---|---|
| gm-AE\|1 | 698.2858 |
| gm-AF\|1 | 716.3116 |
| gm-AI\|1 | 682.3273 |
| gm-AK\|1 | 697.3382 |
| gm-AQ\|1 | 697.3018 |
| gm-AW\|1 | 755.3225 |
| gm-AY\|1 | 732.3065 |
| gm-AEJ\|1 | 870.3706 |
| gm-AEJA\|1 | 941.4077 |
| gm-AEJD\|1 | 985.3975 |
| gm-AEJF\|1 | 1017.439 |
| gm-AEJG\|1 | 927.392 |
| gm-AEJH\|1 | 1007.4295 |
| gm-AEJI\|1 | 983.4546 |
| gm-AEJK\|1 | 998.4655 |
| gm-AEJM\|1 | 1001.4111 |
| gm-AEJN\|1 | 984.4135 |
| gm-AEJT\|1 | 971.4183 |
| gm-AEJV\|1 | 969.439 |
| gm-AEJW\|1 | 1056.4499 |
| gm-AEJY\|1 | 1033.4339 |
| gm-AEJAA\|1 | 1012.4448 |
| gm-AEJAG\|1 | 998.4292 |
| gm-AEJAI\|1 | 1054.4918 |
| gm-AEJAK\|1 | 1069.5027 |
| gm(-Ac)-AEJ\|1 | 828.36008 |
| gm(-Ac)-AEJA\|1 | 899.39722 |
| gm-A\|1 | 569.24331 |

**Table S5. Species, taxids, and genome accessions used for comparative genomics.**

| Species | Taxid | Accession |
|---|---|---|
| *Escherichia coli* | 562 | GCF_000005845.2 |
| *Clostridioides difficile* | 1496 | GCF_018885085.1 |
| *Mycobacterium tuberculosis* | 1773 | GCF_000195955.2 |
| *Enterococcus faecium* | 1352 | GCF_009734005.1 |
| *Sphingomonas paucimobilis* | 13689 | GCF_016027095.1 |
| *Erythrobacter litoralis* | 39960 | GCF_001719165.1 |
| *Caulobacter vibrioides* | 155892 | GCF_000022005.1 |
| *Asticcacaulis biprosthecum* | 76891 | GCF_000204015.1 |
| *Roseobacter denitrificans* | 2434 | GCF_002983865.1 |
| *Rhodobacter sphaeroides* | 1063 | GCF_000021005.1 |
| *Bartonella grahamii* | 33045 | GCF_000022725.1 |
| *Labrys okinawaensis* | 346911 | GCF_002982075.1 |
| *Aquamicrobium aerolatum* | 561088 | GCF_900113935.1 |
| *Mesorhizobium mediterraneum* | 43617 | GCF_002284565.1 |
| *Agrobacterium tumefaciens* | 358 | GCF_003667905.1 |
| *Sinorhizobium meliloti* | 110321 | GCF_007827695.1 |
| *Angulomicrobium tetraedale* | 217068 | GCF_014195655.1 |
| *Hyphomicrobium denitrificans* | 53399 | GCF_000143145.1 |
| *Hirschia baltica* | 2724 | GCF_000023785.1 |
| *Hyphomonas sediminis* | 2866160 | GCF_019679475.1 |
| *Magnetospirillum gryphiswaldense* | 55518 | GCF_000513295.1 |
| *Thalassospira lucentensis* | 168935 | GCF_000421265.1 |
| *Roseomonas gilardii* | 257708 | GCF_001941945.1 |
| *Acidiphilium facilis* | 525 | GCF_000687875.1 |
| *Acidomonas methanolica* | 437 | GCF_004346035.1 |
| *Komagataeibacter xylinus* | 28448 | GCF_004006375.1 |
| *Gluconobacter oxydans* | 442 | GCF_000583855.1 |
| *Gluconobacter frateurii* | 38308 | GCF_002723955.1 |
| *Acetobacter pasteurianus* | 438 | GCF_009914215.2 |
| *Acetobacter tropicalis* | 104102 | GCF_001580945.1 |
| *Acetobacter pomorum* | 65959 | GCF_002738225.1 |
| *Acetobacter aceti* | 435 | GCF_000379545.1 |

**Table S6.** Bacterial strains, plasmids, and oligonucleotides.

| Strains/plasmids/ oligonucleotides | Relevant properties/sequence [a] | Source |
|---|---|---|
| **Strains** | | |
| *Gluconobacter oxydans* | | |
| B58 | *G. oxydans* reference strain (ATCC NRLL B58) | ATCC |
| 621H | *G. oxydans* reference strain (ATCC 621H) | ATCC |
| B58 $ldt_{Go1}$ | B58 with a transposon insertion in Go2094 (GOX2269 in strain 621H); KanR | (1) |
| B58 $ldt_{Go2}$ | B58 with a transposon insertion in Go2227 (GOX1074 in strain 621H); KanR | (1) |
| B58 $ldt_{Go2}$ (pBBR-TetR-Go2227) | B58 $ldt_{Go2}$ mutant complemented; KanR GmR | This work |
| *Escherichia coli* | | |
| BL21(DE3) Lemo | Expression strain | NEB |
| BL21(DE3) (pET-Go2227) | BL21(DE3) Lemo expressing $\mathrm{Ldt}_{\mathrm{Go2}}$ ; AmpR | This work |
| **Plasmids** | | |
| pET2818 | pET28a derivative for recombinant protein expression; AmpR | (2) |
| pET-$\mathrm{Ldt}_{\mathrm{Go2}}$ | pET2818 derivative encoding full length $\mathrm{Ldt}_{\mathrm{Go2}}$; AmpR | This work |
| pBBR1MCS-5 | pBBR1-MCS derivative; GmR | (3) |
| pBBR1MCS-5-$\mathrm{T}_{\mathrm{gdhM}}$-*tetR-mNG* | pBBR1-MCS5 derivative for inducible expression with anhydrotetracycline; GmR | (4) |
| pBBR-TetR-Go2227 | pBBR1-TetR derivative expressing full length $\mathrm{Ldt}_{\mathrm{Go2}}$; GmR | This work |
| **Oligonucleotides** | | |
| SM_0725 | ggctacggtctcccgaagtctcgggccgtctcttgggctt | |
| SM_0726 | ggctacggtctcttccttggattcacttttctctatcactgataggg | |
| SM_0727 | ggctacggtctctaggagatatcatatgcgtgatgtttccagactgac | |
| SM_0728 | ggctacggtctcaaaggtaacggtcttttatccgcaatag | |
| SM_0729 | ggctacggtctcaccttaacgcaaaaaaccccgcttcggcgg | |
| SM_0730 | ggctacggtctctttcgggagcgcctgaagcccgtt | |

[a] KanR, resistant to kanamycin; GmR, resistant to gentamicin; AmpR, resistant to ampicillin


(1) Schmitz, A. M., Pian, B. Medin, S., Reid, M. C., Wu, M., Gazel, E., Barstow, B. (2021) Generation of a *Gluconobacter oxydans* knockout collection for improved extraction of rare earth elements. Nat Commun **12**(1): 6693

(2) Eckert, C., Lecerf, M., Dubost, L., Arthur, M. and Mesnage, S. (2006) Functional analysis of AtlA, the major *N*-acetylglucosaminidase of *Enterococcus faecalis*. *J Bacteriol.* **188**(24):8513-8519

(3) Kovach, M. E., Elzer, P. H., Hill, D. S., Robertson, G. T., Farris, M. A., Roop, R. M., Peterson, K. M. (1995) Four new derivatives of the broad-host-range cloning vector pBBR1MCS, carrying different antibiotic-resistance cassettes. *Gene.* **166**(1):175–6.

(4) Fricke, P. M., Lürkens, M., Hünnefeld, M., Sonntag, C. K., Bott, M., Davari, M. D., Polen, T. (2021) Highly tunable TetR-dependent target gene expression in the acetic acid bacterium *Gluconobacter oxydans*. *Appl Microbiol Biotechnol.* **105**(18):6835-6852.
